# Using a sequential sampling algorithm to apply the niche-neutral model to species occurrence patterns

**DOI:** 10.1101/2024.11.07.622409

**Authors:** Nadiah P. Kristensen, Yong Chee Keita Sin, Hyee Shynn Lim, Frank E. Rheindt, Ryan A. Chisholm

## Abstract

**Aim:** Species occurrence patterns are typically analysed using data-randomisation approaches, which reveal when observed patterns deviate from random expectation, but give little insight why. When non-randomness is detected, the analysis reaches a dead end. Mechanistic models, such as neutral models, offer an alternative: when their predictions fail to match data, the specific nature of each mismatch can implicate candidate mechanisms, turning null-model rejection into a diagnostic process. However, mechanistic models can be computationally expensive. Here, we use an efficient method to simulate such models and explore possible mechanisms governing the occurrence patterns of birds on islands.

**Location:** Riau archipelago, Indonesia.

**Taxon:** Birds.

**Methods:** We used species richness and island–area data to fit a niche–neutral model, where species obey neutral dynamics within non-overlapping discrete niches. We used a sequential sampling algorithm that can efficiently sample presence–absence matrices under the niche-neutral model, and used mismatches to identify which mechanisms were potentially important to occurrence patterns. In particular, we compared model to observed data using standardised effect sizes on segregation (C-score) and nestedness (NODF) metrics.

**Results:** Birds were more segregated and less nested than expected from both data randomisation and the niche–neutral model. Further, while the niche–neutral model reproduced the mean relationship between island size and species richness, it could not produce sufficient variability to account for richness variation across islands. However, while the niche–neutral model was rejected as a null, it was possible to reproduce the species-occurrence patterns by allowing niche diversity and per-capita immigration rate to vary across islands, which increased segregation and decreased nestedness, respectively.

**Main conclusion:** While the species-area relationship could be explained by a model with constant per-capita immigration rates and number of niches across islands, inter-island heterogeneity was needed to explain species-occurrence patterns. Unlike data randomisation, which would have identified the patterns as non-random but offered no further insight, the mechanistic approach identified habitat diversity and immigration-rate variation as candidate mechanisms, demonstrating the diagnostic value of using niche–neutral models as an exploratory framework. The sequential sampling algorithm allowed us to explore different scenarios efficiently and may be useful for identifying potential mechanisms structuring patterns in other systems.

## 1. Introduction

What determines the pattern of species occurrence across patches in a landscape? One method for identifying patterns of potential ecological interest is null-model analysis using randomised data (Gotelli, 2000; Zhang, 2020). One first computes a metric that summarises an occurrence pattern, then one compares the observed value to the distribution obtained when the data are randomised in various ways (Ulrich and Gotelli, 2007). For example, the ‘fixed–fixed’ randomisation swaps species presences and absences (1s and 0s in the presence–absence matrix) while holding fixed the number of sites where each species is observed (row sums) and the number of species observed at each site (column sums) (Strona et al., 2014). If the observed metric deviates significantly from the distribution in randomised data, then qualitative inferences can be drawn about the underlying mechanisms (e.g., Kohli et al., 2018; Meyer and Kalko, 2008). For example, if the observed co-occurrences between species occupying the same niche are more segregated than in randomised data, i.e., the “checkerboard pattern” (Connor and Simberloff, 1979; Gotelli, 2000), then this may indicate strong interspecific competition (Diamond, 1975).

Data randomisation tests are versatile (non-parametric and distribution free) and impose minimal data requirements, i.e., no information other than the presence–absence matrix itself is required; however, its use as a null model lacks a mechanistic foundation. In statistical hypothesis testing, the null should instead represent the same system with the ecological mechanisms of interest excluded (Bausman, 2018; Molina and Stone, 2020; Zhang, 2020). While data randomisation is useful for challenging intuitions about how unusual an observed pattern really is (Zhang, 2020), ecologists could benefit from a mechanistic model framework that allows more rigorous quantitative testing of the mechanisms driving species occurrence patterns.

Crucially, when data randomisation reveals a significant deviation from the null, the analysis reaches a dead end: one knows that the pattern is non-random, but no framework for investigating which mechanisms produced it. A mechanistic model, by contrast, converts that rejection into a starting point: the specific ways in which model predictions fail to match observations can generate testable hypotheses about missing mechanisms.

One possible source of mechanistic null models is neutral theory (Bell, 2005; Molina and Stone, 2020; Munoz and Huneman, 2016; Rosindell et al., 2012). Neutral models assume an equivalence between species and focus on predicting broad statistical patterns of biodiversity (e.g., species-abundance distribution (Missa et al., 2016; Ofiţeru et al., 2010; Rosindell et al., 2012; Volkov et al., 2003). In particular, the spatially implicit neutral model (Hubbell, 2001) takes a macroecological perspective (e.g., MacArthur and Wilson, 1967) in which communities are treated as subsets of a regional community and are formed by broad-scale stochastic processes: immigration and stochastic extinction due to drift (Munoz and Huneman, 2016). The advantages of the spatially implicit neutral model are that it is mechanistic (Alonso et al., 2006) and analytically and computationally tractable (Rosindell et al., 2012) to a degree that may make null-model testing feasible. In this study, we focus on niche–neutral models, which incorporate some degree of niche mechanisms into the mathematical framework of neutral theory (e.g., Chisholm and Pacala, 2010, 2011).

A neutral model (e.g., Gravel et al., 2006; Munoz et al., 2018) could serve as a baseline, representing expectations based on established processes, to help distinguish the relative contributions of neutral and other factors in producing the observed patterns (Bausman, 2018). For example, one may start with a neutral model, and if the fit is unsatisfactory, add features that are expected to be important and test whether they improve the fit. Although the model used will necessarily be incomplete, it may still provide a relevant explanation for biodiversity in terms of the main processes involved (Odenbaugh, in press), for example, if co-occurrence patterns are mainly structured by dispersal and stochasticity in the birth, death, immigration, and speciation processes (cf., Bell, 2001). Further, the ways in which the model predictions differ from observed data may reveal important mechanisms missing from the model (Marquet et al., 2014; Matthews and Whittaker, 2014; Rosindell et al., 2012), and may provide a backdrop against which more complex mechanisms can be tested (Blanchard et al., 2020; Munoz and Huneman, 2016; Volkov et al., 2003).

One barrier to using a mechanistic model as a null model is that model parameterisation requires more data (e.g., Gotelli and McGill, 2006). Data randomisation requires only the data itself, i.e., the presence-absence matrices, whereas mechanistic macroecological models typically require estimates of other parameters, such as the immigration rate and metacommunity diversity (Adler, 2004; Chisholm et al., 2016; Wootton, 2005). To estimate these parameters, species-abundance data is typically needed (Beeravolu et al., 2009). Although a few studies have explored how co-occurrence and nestedness patterns under neutral models differ from random in simulations (Bell, 2005; Gotelli and McGill, 2006; Ulrich, 2004; Ulrich et al., 2017), no study to date has used a neutral model as an alternative to the traditional data-randomised presence-absence matrix when analysing these patterns in empirical data.

Here, we explore the potential of a niche–neutral model (Chisholm et al., 2016) to overcome the parameterisation problem and provide a baseline for exploring when species occurrence patterns can be explained by just a few simple mechanisms. It has the advantage that species-abundance data is not needed, only presence-absence and island-area data.

Our approach treats the niche–neutral model not as a definitive description of the system, but as a diagnostic instrument. We begin with the simplest version of the model, identify specific ways in which its predictions fail to match the data, and then ask which ecologically plausible modifications resolve each mismatch. The value of the exercise lies not in the final parameterisation — which is deliberately overfitted — but in what the pattern of mismatches and their resolution reveals about which mechanisms are likely to matter.

Our application is to bird species on islands in an archipelago in Southeast Asia (Sin et al., 2022). The niche–neutral model used is an extension of Hubbell’s spatially implicit neutral model. Each patch within the model comprises the same set of distinct equal-sized non-overlapping habitats or niches, and species obey neutral dynamics within each niche (Chisholm et al., 2016). To efficiently sample presence–absence data from this model, we extend an existing sequential sampling scheme (Etienne, 2007). We found that the occurrence patterns were inconsistent with the null distributions generated by the most basic version of this model, and that inter-island variation in immigration rate and niche composition were required to improve fidelity to the data. The approach thus leads to better mechanistic understanding of the patterns observed (Zhang, 2020).

## 2. Methods

### 2.1. Case study

We investigated a presence–absence matrix obtained from a comprehensive survey of birds on 23 islands in the Riau region, Indonesia, a subset of data previously published by Sin et al. (2022). The dataset included a substantial number of small islands (*<*10 km^2^) to capture the small-island effect as well as the classic increase phase of the species–area curve (Chisholm and Fung, 2022). Surveys were conducted by traversing the islands (Sodhi et al., 2010) and excluded species that were introduced, nocturnal, aquatic, pelagic, coastal, and those not breeding on the islands (i.e., migratory species and flyby birds of prey). Species inventories were available for two islands to supplement observations (Lingga: Sutari (2004) and Gibson-Hill (1952); Durian: Sieber (1926)). To produce the dataset including the 23 islands we used, Sin et al. (2022) assessed inventory completeness using species accumulation curves (Oksanen et al., 2019), and 7 islands with poor sampling were excluded (Weigelt and Kreft, 2013).

### 2.2. Niche–neutral model

The niche–neutral model (Chisholm et al., 2016; Chisholm and Pacala, 2010, 2011) partitions the local island community of *J* individuals into *n*^∗^ distinct non-overlapping niches whose dynamics are completely independent. Within each niche, the community dynamics are neutral. At each time step, an individual is chosen at random to die, and it is replaced by either: an immigrant from the metacommunity, with probability *m*; or the offspring of another randomly chosen individual that shares its niche, with probability 1 − *m*.

The metacommunity is also divided into *n*^∗^ niches, and the total number of individuals is equal to *J*_*M*_ . The metacommunity undergoes a similar birth–death dynamics to the island community, with the key difference that species input into the metacommunity occurs via point speciation, with per-capita probability *ν*, rather than via immigration. The island community dynamics operate on a much faster timescale than the metacommunity such that the metacommunity can be assumed static on timescales relevant to the local community. A composite parameter, named the fundamental biodiversity number *θ* = (*J*_*M*_ (1 − *ν*))*/ν*, characterises the diversity of the metacommunity at equilibrium (Hubbell, 2001). It is assumed that *θ* is equal across niches.

Studies of island archipelagos provided the first empirical support for the niche–neutral model (Chisholm et al., 2016), and subsequent experimental work with benthic invertebrates on intertidal sea-walls has confirmed its general processes (Loke and Chisholm, 2023). In archipelagos, the model explains a key relationship observed between island area and species richness (Chisholm et al., 2016) called the small-island effect (e.g., Lomolino and Weiser, 2001). In the model, equilibrium species richness generally increases with island area, driven by the effect of area on the immigration–extinction stochastic equilibrium, as in classic island biogeography theory (MacArthur and Wilson, 1967). But among small islands, the model departs from classic island biogeography theory by exhibiting a nearly flat phase that is consistent with the small-island effect seen in archipelagic data. In the small-island phase, equilibrium species richness is not driven by immigration–extinction dynamics but is instead roughly equal to the number of niches (Chisholm et al., 2016).

To compare the model to data, four parameters must be estimated (Chisholm et al., 2016): (1) the number of niches, *n*^∗^; (2) the per-capita immigration rate, *m*; (3) the fundamental biodiversity parameter, *θ* (Hubbell, 2001); and (4) the density of individuals per unit area, *ρ*. The last of these can often be obtained from independent density estimates for the species group concerned, but the other three parameters must typically be fitted to the archipelago richness data via least squares or a related method. Our procedure was to find the best-fit values of *m* and *θ* for each value of *n*^∗^, and then choose from among the *n*^∗^ the (*n*^∗^, *m, θ*) triple that gave the highest *R*^2^ value (details in SI B).

In the simplest version of the niche–neutral model, the homogeneous niche-neutral model, the niches are all assumed to have equal size, all islands are assumed to have the same number of niches, *n*^∗^, and all niches are available on every island.

### 2.3. Sequential sampling scheme

To efficiently simulate presence–absence data from the niche–neutral model, we extended the work of Etienne (2005, 2007) to create a sequential sampling scheme equivalent to the niche–neutral model (see also Munoz et al., 2007). The sampling scheme generates a random sample of the species identity of every individual in the archipelago, and from this a synthetic presence–absence matrix can be derived. Although determining the species identity of every individual is more computationally expensive than data randomisation, it is more efficient than forward-time simulation — the scheme samples only present-day individuals plus their immigrant ancestors rather than every individual who was historically present.

Our sequential sampling scheme determines each individual’s species identity by sampling events in the coalescence tree (Etienne and Olff, 2004; Rosindell et al., 2008). It assumes that speciations do not occur at the island-community level during community dynamics, but only on the metacommunity level. In the context of an island community, at each step backwards in time, a lineage may either coalesce with another lineage of the community, which indicates that all individuals in both lineages share a common ancestor in that community and thus belong to the same species, or it may terminate in an immigrant ancestor. The immigrant is drawn from the metacommunity and can belong to a species already present in the community or to a new species. In the context of the metacommunity, immigrant lineages may either coalesce with another lineage or terminate in a point-speciation event. Thus, the species identity of all individuals in a lineage is determined when the lineage is traced back to a speciation event. The niche–neutral model assumes neutral dynamics within each niche, which is equivalent to the coalescence model (Rosindell et al., 2008), and the coalescence model is in turn equivalent to a sequential sampling scheme (Branson, 1991; Kingman, 1982; Wakeley, 2009). Thus, our sequential sampling scheme produces samples that are statistically identical to those of conventional forwards-time simulations Kingman (1982) but orders-of-magnitude more efficiently.

Our sampling scheme builds on three prior schemes, which we will discuss in order of increasing algorithm complexity: (1) the metacommunity, (2) the island community, and (3) multiple island communities.

To sample from the metacommunity, we model the coalescence process by drawing each individual’s species-identity-determining common ancestor one at a time. This can be physically represented by an urn model (Fig. 1a). The process is governed by the fundamental biodiversity number *θ* (Hubbell, 2001), which increases with community size and speciation rate and determines overall species diversity (cf., Ewens, 1972). Each draw corresponds to an event in the coalescence tree, either a coalescence (Eq. 1a) or a point-speciation event (Eq. 1b). For the *j*th individual drawn, the probability of it belonging to a previously sampled species is

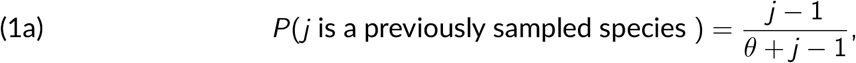

while the probability of it belonging to a new species is

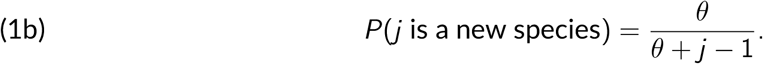

**Figure 1.**
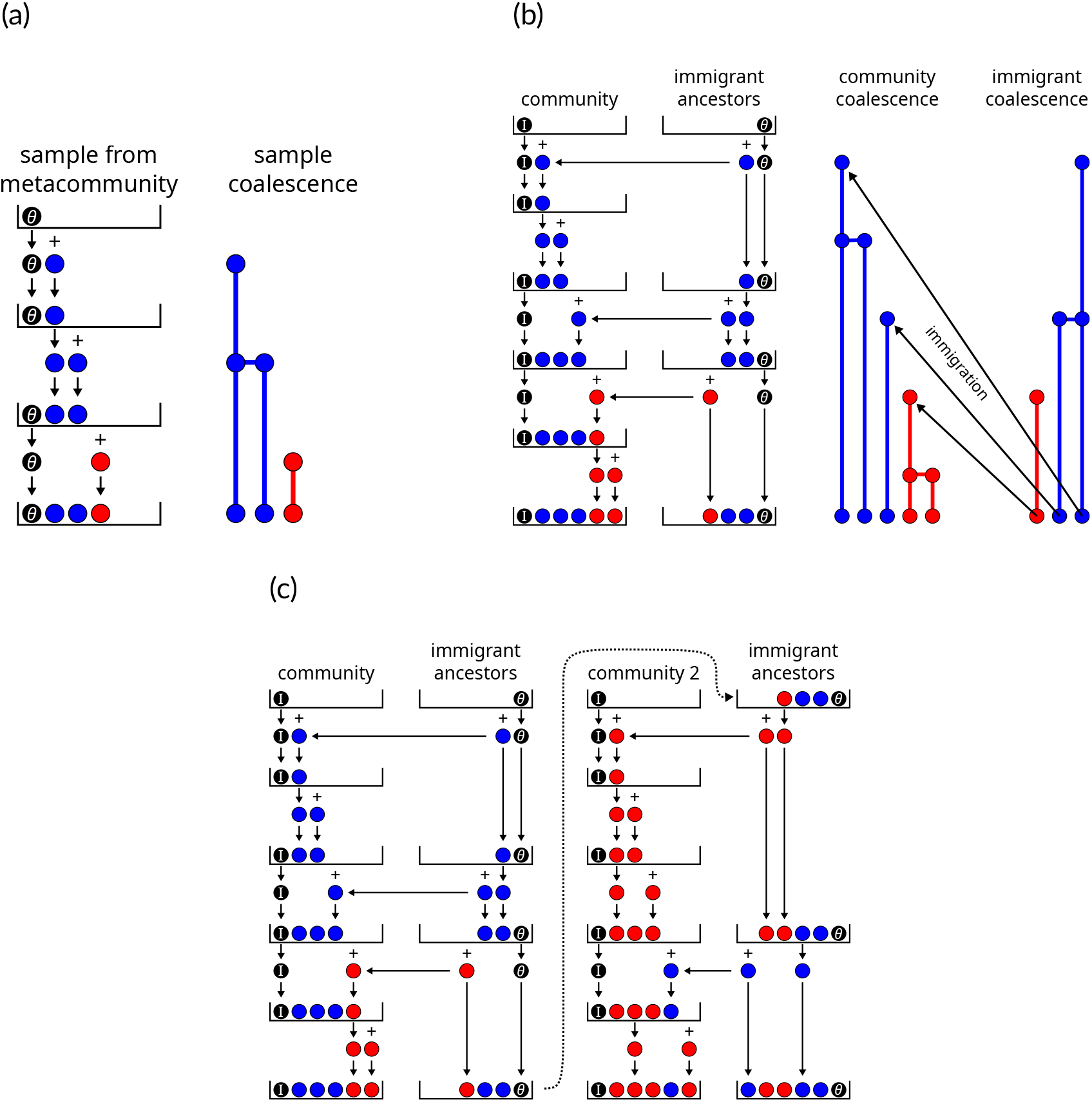
Examples of coalescence models represented by urn models generating samples from (a) a metacommunity, (b) a local community, and (c) a collection of communities sharing a metacommunity (e.g., archipelago). Each coloured ball represents an individual, its colour indicates its species identity, and all coloured balls have equal weight 1 reflecting their equivalence in the neutral model. Black balls represent an immigration with weight *I* and a speciation with weight *θ*. **(a)** The black speciation ball *θ* is placed in the urn. At each step, a ball is drawn and returned, and a new ball is added depending on the colour of the ball drawn. If a coloured ball is drawn, that represents branching in the coalescence tree (Eq. 1a), and a ball of the same colour is added. If the black ball is drawn, that represents speciation (Eq. 1b), and a ball of a new colour is added. **(b)** A black immigration ball *I* is placed in the ‘community’ urn and speciation ball *θ* is placed into the ‘immigrant ancestors’ urn. At each step, the ‘community’ urn is sampled first. If a coloured ball is drawn, a ball of the same colour is added to the ‘community’ urn only. If the immigration ball is drawn, a draw from the ‘immigrant ancestors’ urn determines the colour of the ball to be added to both urns: a coloured ball adds the same colour to both urns, the speciation ball adds a new colour to both urns. **(c)** There is a single ‘immigrant ancestors’ urn and a separate ‘community’ urn for each local community. ‘Community’ urns are filled sequentially according to the same rules as (b). Once a community is filled, the ‘immigrant ancestors’ urn and its current state is passed to the next community.

Eq. 1 is also known as the ‘species generator’ (Chapter 9, Hubbell, 2001; Rosindell et al., 2011), and it was derived first in the genetics literature from the analogous neutral genetic process (Lemma 2.1, Donnelly (1986); see also: Ewens (1972), Kingman (1982), and Moran (1958)).

To simulate an island community, a model with two urns is used: one tracks coalescence events on the island, while the other tracks immigrants from the metacommunity (Etienne, 2005) (Fig. 1b). On the island, new species arrive by immigration instead of speciation, which is governed by an immigration rate 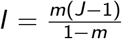 analogous to the role of *θ* previously. These immigrants are treated as a random sample from the metacommunity and modelled as described above (i.e., Fig. 1a). This simplifies reality by approximating the distribution of immigrant identities over time by a sample drawn at a single point in time (which is why the arrows representing immigration in Fig. 1b all start at the base of the immigrant-coalescence tree).

To simulate multiple islands, each island is represented by one urn, while a shared immigrant-ancestors urn serves as the common source for all islands (Etienne (2007); Fig. 1c). This setup aligns with the niche–neutral model’s assumption that islands are connected by immigration to the same metacommunity, but not each other (a necessary mathematical simplification). Although the sequence of draws from the ancestor urn corresponds to a temporal sequence of events in the coalescence tree, for a given final outcome, the probability of every temporal sequence is the same. As a result, Etienne (2007) was able to streamline the scheme by sampling the islands sequentially without altering the probability distribution of outcomes (cf., Munoz et al., 2007).

To simulate the complete niche–neutral model, the multiple-island simulations (see Fig. 1c) are replicated for each niche 1, …, *n*^∗^ independently (see SI A). Our sequential sampling scheme also allows us to extend the scenario presented by Chisholm et al. (2016) by allowing the number of niches and the immigration rates to vary between islands (heterogeneous models below).

### 2.4. Species-occurrence patterns: co-occurrence and nestedness

We used two metrics to quantify species-occurrence patterns in both our data and models, which we chose because they are widely used and well-established in the literature: C-score to quantify co-occurrence (Stone and Roberts, 1990) and NODF to quantify nestedness (Almeida-Neto et al., 2008). We calculated the C-score using the `EcoSimR`package for `R`(Gotelli et al., 2015), and we calculated NODF using code adapted from Strona et al. (2018). In arranging the presence–absence matrix, species rows were sorted by decreasing fill, while columns representing islands were ordered by island size, which reflected the anticipated effect of island area on species richness.

To allow comparison between real and simulated data, we also computed each metric’s standardised effect size (SES) using the fixed-fixed algorithm (Fig. 2e). When comparing across studies, it is generally recommended to use SES rather than raw metric scores (Gotelli and McCabe, 2002). Furthermore, Ulrich et al. (2018) also found that SES scores can covary with total species richness in a nonlinear way, and the only approach that allowed meaningful comparisons was when SES scores were calculated using the fixed-fixed algorithm. The fixed-fixed algorithm also has favourable statistical properties when used with C-score (Gotelli, 2000) and other nestedness metrics (Ulrich and Gotelli, 2007) (however, see Molina and Stone (2020) for a conceptual critique of those favourability criteria). The downside is that calculating fixed–fixed SES scores is computationally expensive.

**Figure 2.**
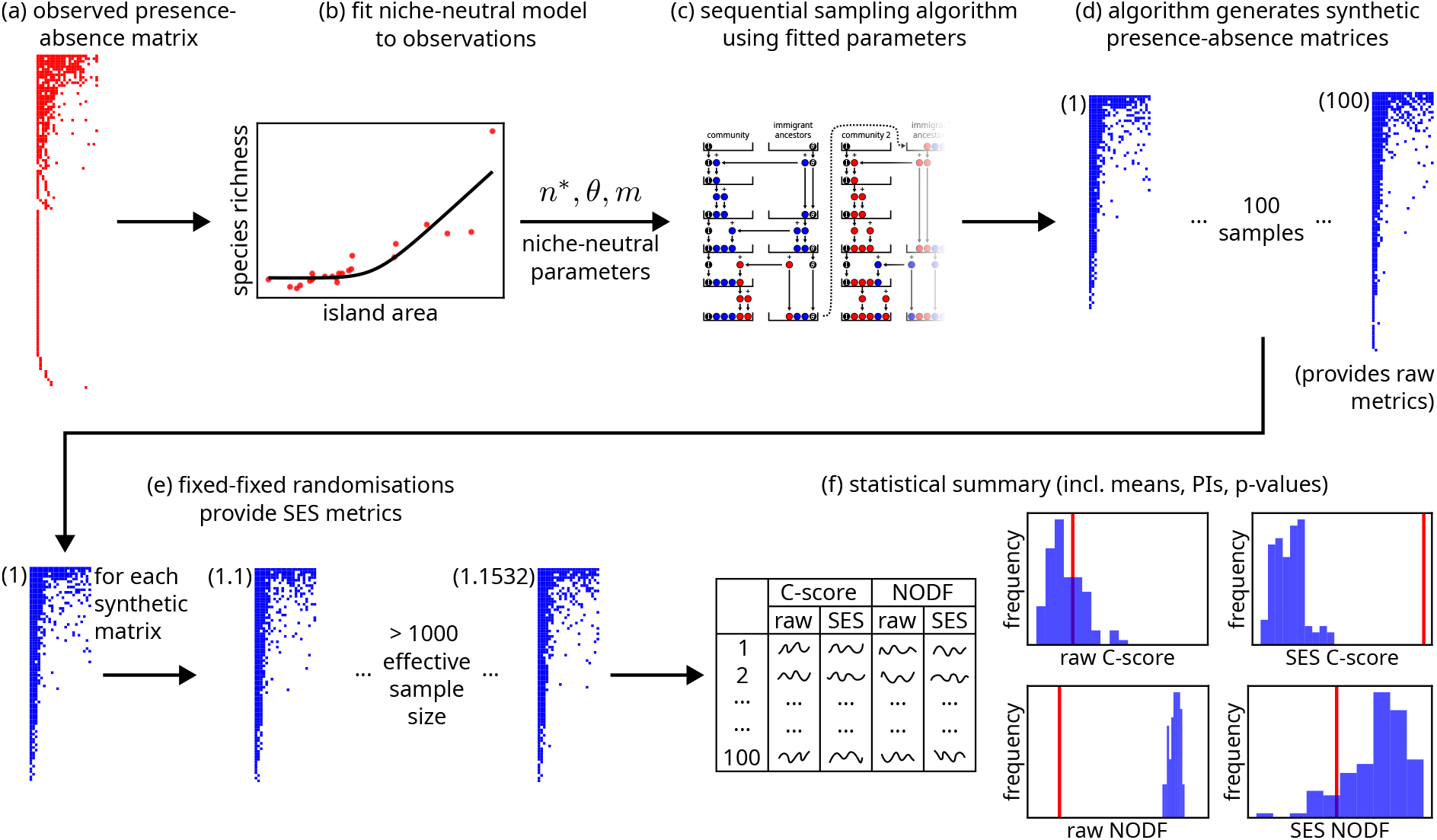
Overview of the method used to summarise the homogeneous niche-neutral model’s predictions and compare them to the observed data.

The fixed–fixed randomisation was performed using the curveball algorithm (Strona et al., 2014) implemented in the `EcoSimR`package for `R`(Gotelli et al., 2015); however, we added two additional steps to account for autocorrelation between samples during randomisation. First, we visually inspected the traceplots (i.e., plots of the metric value versus iteration number) to verify that the burn-in length was sufficient. Second, we calculated the effective sample size using the `CODA`package for `R`(Plummer et al., 2006) to ensure that metrics were calculated from an effective sample size of at least 1000 samples.

The SES scores were calculated as follows. Let *V*_raw_ denote the raw metric value (C-score or NODF) calculated from a presence–absence matrix, which may originate from the survey data or synthetic data sampled from the niche–neutral model; let 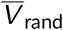 denote the mean value when the matrix is randomised according to the fixed–fixed algorithm; and let *σ*_rand_ denote its standard deviation. Then

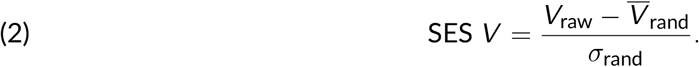

Note that the randomisation used to calculate the SES scores here does not represent a “null” hypothesis, but rather it serves to scale the metrics so their values can be more meaningfully compared across data and model samples (Fig. 2f).

### 2.5. Can niche–neutral mechanisms reproduce the observations?

We explored the niche–neutral model in two stages. In the first stage, we fitted a homogeneous model to the species-area data and compared the species-occurrence patterns in the synthetic presence-absence matrices to the observed patterns (Fig. 2). The homogeneous model was rejected as a null model. However, the homogeneity assumption is unrealistic and not a necessary assumption for niche–neutral processes.

In the second stage, we therefore relaxed the homogeneity assumption to determine whether any heterogeneous niche–neutral model could reproduce the observed patterns. Having established that the homogeneous model was rejected, we used the specific nature of its mismatches to generate hypotheses about which mechanisms were missing. The goal of the following analysis was not to arrive at a well-constrained fitted model, but to verify that identifiable, ecological plausible modifications to the niche–neutral framework could account for each discrepancy. We first verified which niche–neutral mechanisms influenced nestedness and co-occurrence, and then we combined them through manual calibration, as a proof-of-principle demonstration.

The secondary analysis serves as proof-of-principle that purely niche–neutral processes can produce the observed patterns; however, two caveats apply. First, the mechanisms we explored were necessarily restricted to those available in the model; for example, an exploration of trait-based environmental filtering would require an alternative model such as ecolottery (Munoz et al., 2018). Second, allowing the parameters to vary between islands greatly increased the degrees of freedom, allowing the model to be overfitted. Therefore, the goal is not to infer specific parameter values; rather, this analysis demonstrates that niche–neutral processes can generate the observed patterns, and that a mechanistic approach can provide a pathway for model refinement when initial predictions are rejected.

#### 2.5.1. Homogeneous niche-neutral model

The homogeneous model assumes that all islands have the same number of equal-sized niches *n*^∗^ and per-capita immigration parameter *m*. To parameterise the model, we approximated the density of birds as *ρ* = 1700 individuals/km^2^, which was chosen as a compromise between values reported in the literature: 1259 individuals/km^2^ for resident birds in old secondary forests (Castelletta et al., 2005), and 2115 individuals/km^2^ average density for primary rainforest habitats (Sheldon and Styring, 2011). The other three model parameters — number of niches *n*^∗^, immigration rate *m*, and fundamental biodiversity number *θ* — were fitted to the island species–area data using the methods described in Chisholm et al. (2016) (Fig. 2b, details in SI B).

The fitted niche–neutral model parameter values were input to the sequential sampling scheme (Fig. 2c), from which synthetic presence–absence matrices were derived (Fig. 2d) and their species-occurrence metrics compared to observed data (Fig. 2d–f).

#### 2.5.2. Identifying candidate mechanisms using heterogeneous niche-neutral models

We first verified which niche–neutral mechanisms influenced nestedness and co-occurrence, and then we combined them to find a heterogenous model that could reproduce the observed patterns.

We hypothesised that nestedness could be decreased by tuning the immigration rate and number of niches so the model correctly predicted island richnesses, and that segregation could be increased by allowing niche diversity to increase with island area.

Nestedness is largely determined by relative species richnesses among islands: in a nested system, species on less species-rich islands should form a subset of those on more species-rich islands. Recall that the columns of the presence-absence matrix were ordered by island size, anticipating the effect of area on species richness, with the intended effect of maximising NODF nestedness. In the niche-neutral model, once the island area is fixed, species richness is governed by the number of niches *n*^∗^ on small islands and the immigration rate *m* on larger islands. Tuning these parameters so that the model correctly reproduces the observed island-richness pattern is therefore an obvious first step towards addressing any nestedness mismatch.

A neutral model alone will not produce a checkerboard species pattern beyond random chance; however, a species-segregated pattern can be produced by a niche–neutral model through differential habitat preferences (Connor et al., 2013) if some niches are available on some islands but not others. We expect this to be true in general, but particularly because large islands contain key habitats (e.g., rainforests) that are not available on small islands (Davidar et al., 2001; Kohn and Walsh, 1994; Ricklefs and Lovette, 1999).

To verify the nestedness mechanism, we used the *θ* and *n*^∗^ parameter values from the homogeneous model fit and tuned the value of *m*_*i*_ for each island *i* to match observed species richness. This allowed the model to be overfitted because there are more parameters than data points; however, this overparameterisation was necessary to exactly reproduce each island’s observed richness.

To verify the segregation mechanism, we simulated synthetic archipelagos where niche diversity increased with island size. The synthetic archipelago contained 16 islands with areas distributed evenly on a log scale (carrying capacity range *J* = 10^2^ to 10^5^) and parameter values chosen to capture the small-island effect (i.e., *A*_min_ *< A*_crit_ *< A*_max_), and we considered two main scenarios for how niche diversity increased with island size. In Scenario 1, the four largest islands in the archipelago have eight additional niches, with all niches still equal in area. In Scenario 2, we split the ten niches into five coastal and five interior types, and assumed that the width of coastal niches around the perimeter of the islands (assumed circular) remained constant across the archipelago (details in SI D). This reflects the notion that, as island size increases, the area covered by coastal habitat types will grow more slowly than inland types. Both scenarios were compared to a baseline model, where we kept *m* and *n*^∗^ constant across islands, maintaining the homogeneity assumption.

Having verified the two hypothesised mechanisms, we then combined them through manual calibration to find a model that produced closer agreement with the survey data. Again, readers should note this allows the model to be overfitted, and our analysis should be viewed as simply proof-of-principle that a niche–neutral model can be refined in this way to reproduce the data.

## 3. Results

### 3.1. The homogeneous niche–neutral model versus observed data

The homogeneous niche–neutral model provided a good fit to the species–area relationship by capturing the small-island effect (*R*^2^ = 0.81, Fig. 3a); however, the model did not reproduce other aspects of the data well (Table 1 “Homogeneous”; see also SI C). The total archipelagic richness predicted by the model was too low (Fig. 3b), and when the sequential sampling scheme was used to explore the variation in the model, most island richness values fell outside of the 95th percentile of samples (16 of 23, Fig. 3a).

**Table 1.** Species-occurrence metrics observed versus predicted from the niche–neutral models. In the fitted model, both niche diversity and immigration rates were varied across islands as proof-of-principle that the data could be consistent with niche–neutral processes. Note that the SES score is the preferred metric for detecting occurrence patterns (Eq. 2) and to compare the observations to the niche-neutral models.

| Metric | Observed | Niche-neutral model predictions [95-percentile](lower-tail prob.) |  |  |  |
| --- | --- | --- | --- | --- | --- |
|  |  | Homogeneous |  | Fitted |  |
| spp. richness | 125 | 83.3 | [69, 99] ($p < 0.01$) | (tuned to match observed) | |
| raw C-score | 1.28 | 1.19 | [0.835, 1.81] ($p = 0.33$) | 1.64 | [1.28, 2.07] ($p = 0.03$) |
| SES C-score | 6.60 | -0.476 | [-1.76, 1.40] ($p < 0.01$) | 6.16 | [4.02, 8.80] ($p = 0.37$) |
| raw NODF | 57.2 | 82.5 | [80.2, 84.2] ($p < 0.01$) | 60.3 | [55.4, 65.5] ($p = 0.13$) |
| SES NODF | -2.67 | -0.932 | [-4.24, 0.977] ($p = 0.13$) | -4.39 | [-7.78, -1.49] ($p = 0.13$) |

**Figure 3.**
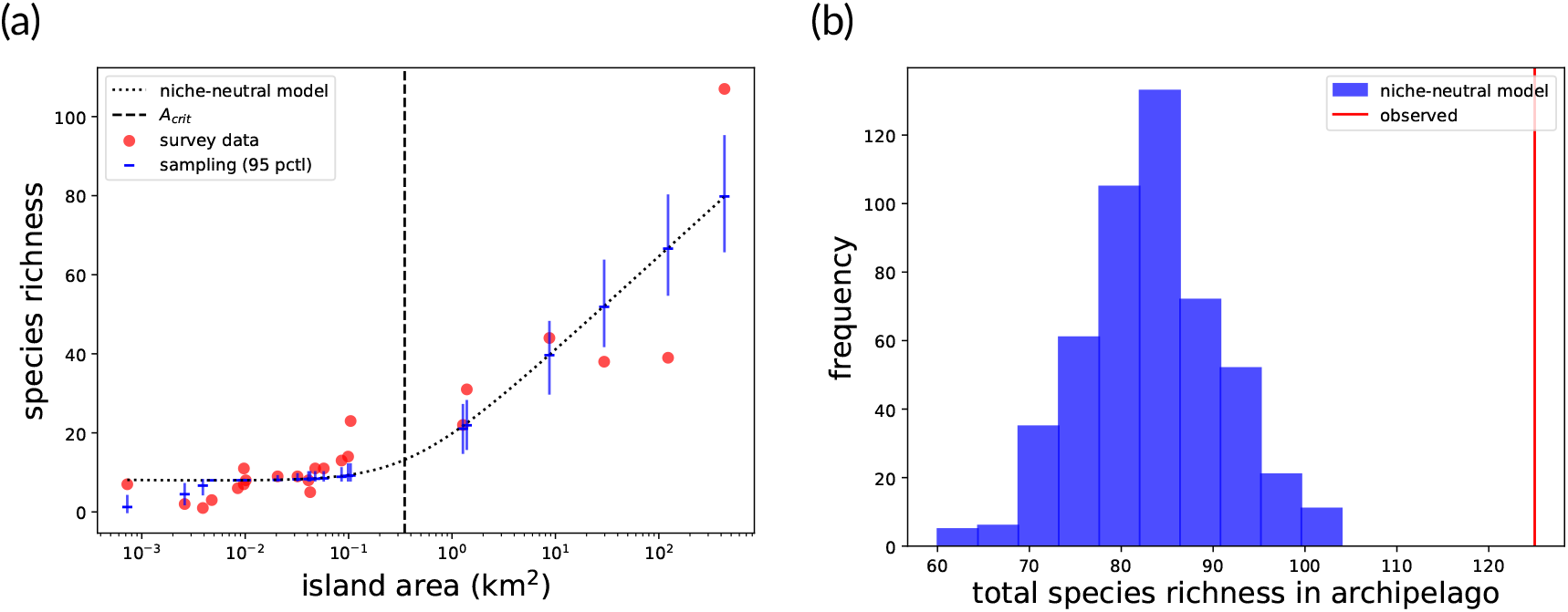
A comparison between the survey data and the predictions of the niche– neutral model. (a) The SAR, best-fitting niche–neutral model predictions (solid line), and 95 percentiles drawn from the sequential sampling scheme. (b) The total archipelago richness (line) compared to the distribution of total richnesses from the sequential sampling scheme. The best-fitting niche–neutral model parameters were: *n*^∗^ = 8, *θ* = 10.3, *m* = 3.6 × 10^−3^, *R*^2^ = 0.81, resulting in *A*_crit_ = 0.35 km^2^.

The raw C-score for the data was within the range of values obtained from the model (data raw C-score 1.28; model raw C-score 1.19, 95% prediction interval (PI) [0.835, 1.81], *p* = 0.33) (Fig. 4a). However, the SES C-score for the data was larger than the model values, indicating segregated patterns in the data (data SES C-score 6.60; model SES C-score −0.476, 95% PI [−1.76, 1.40], *p <* 0.01) (Fig. 4b).

**Figure 4.**
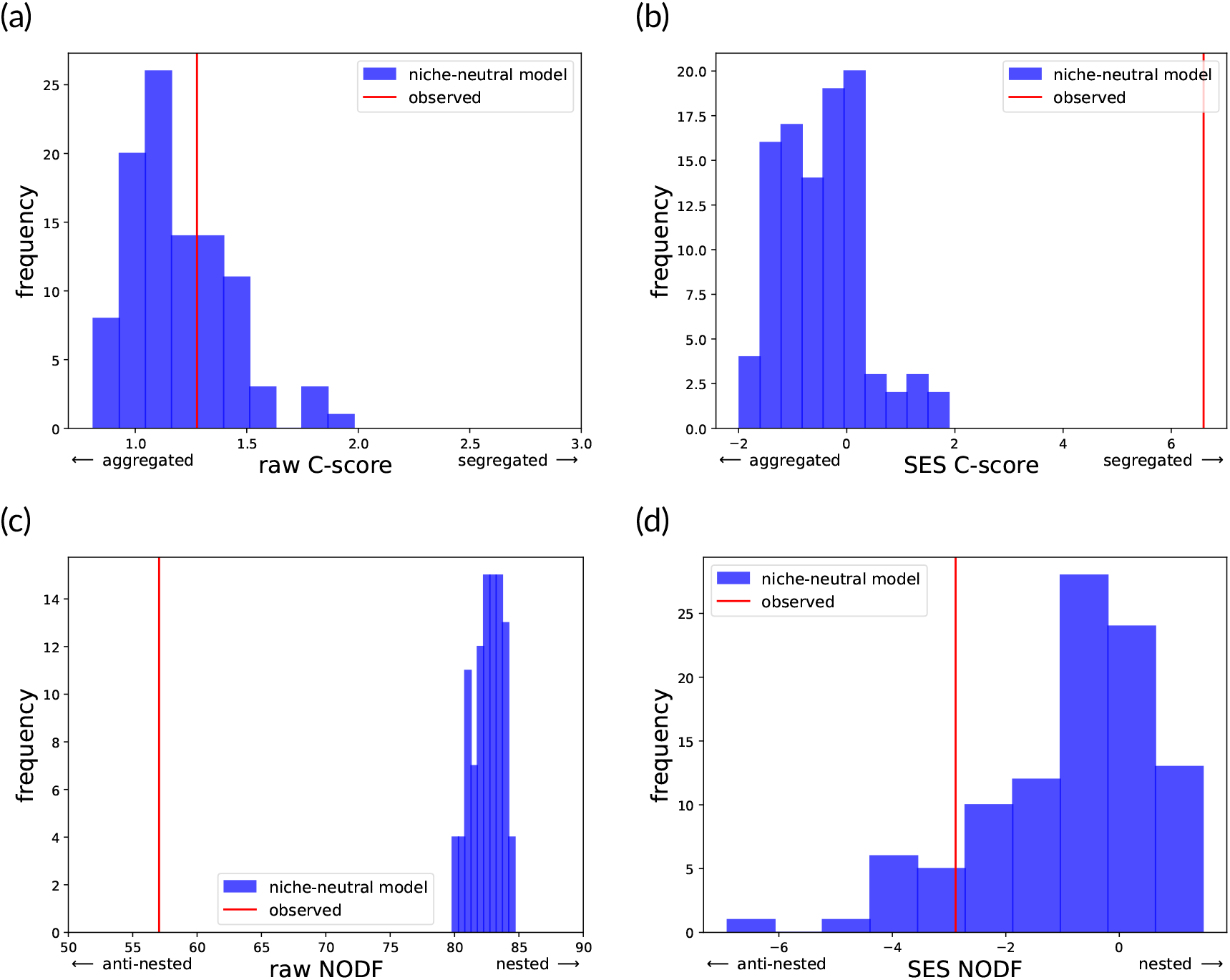
Frequency histograms comparing the distribution of characteristics of the ho-mogeneneous niche–neutral model (blue; 100 draws Fig. 2c-d) to the observed survey data (red): (a) raw C-score, (b) SES C-score, (c) raw NODF, (d) SES NODF. The SES is calculated in comparison to fixed–fixed randomised matrices, which were generated using the sim9 algorithm with an effective sample size of at least 1000 (Fig. 2e).

The niche–neutral model predicted higher NODF scores than observed in the survey data. The disagreement between model and data was stronger for the raw NODF scores (data raw NODF 57.2; model raw NODF 82.5, 95% PI [80.2, 84.2], *p <* 0.01; Fig. 4c) than in the standardised scores (data SES NODF −2.67; model SES NODF −0.932, 95% PI [−4.24, 0.977], *p* = 0.13; Fig. 4d).

A traditional data-randomisation approach produced results that were broadly consistent with the comparison to the niche-neutral model above (Table 2). Specifically, the survey data was more segregated and anti-nested than the niche-neutral model, and the data-randomisation approach also found that the survey data was significantly segregated and anti-nested compared to the fixed-fixed randomisations.

**Table 2.** Null-model analysis using randomised data for bird occurrence in Riau archipelago, including standardised effect sizes (SES) and associated lower-tail probabilities (*p*). Note that the SES score is the preferred metric for detecting occurrence patterns (Eq. 2), and it is used to compare the observations to the niche-neutral model in Table 1.

| Metric | Observed | Mean randomised |  | Interpretation |
| --- | --- | --- | --- | --- |
| C-score | 1.28 | 1.05 | (SES = 6.60; $p < 0.0003$) | Species are segregated across islands. |
| NODF | 57.2 | 59.0 | (SES = -2.67; $p = 0.01$) | Archipelago assemblage is anti-nested. |

While the data-randomisation analysis ends here with the conclusion that the patterns are non-random, the niche—neutral model provides a framework for asking why. The specific mismatches between the homogeneous model and the data (excess segregation, insufficient nestedness, too little richness variability) each suggest candidate mechanisms, which we investigate in the following subsections.

### 3.2. Identifying candidate mechanisms using heterogeneous niche–neutral models

The homogeneous niche–neutral model above fits poorly by under-predicting total species richness, under-predicting segregation, over-predicting nestedness, and failing to capture variation in individual island richness. However, while the homogeneous model is rejected as a null model, a heterogeneous model could plausibly match the observations, and our mechanistic approach provides a framework for exploring that.

The following subsections use the fitted heterogeneous models as diagnostic tools rather than as candidate descriptions of the system. In each case, the question is not whether the modified model is the correct model, but whether a specific, ecologically motivated modification resolves a specific mismatch, thereby implicating the corresponding mechanism.

In subsection 3.2.1, we investigate whether allowing per-capita immigration to vary across islands — in a way that explains the island-specific species-richness — can decrease nestedness. In subsection 3.2.2, we investigate whether incorporating the general empirical relationship between habitat diversity and island size can increase segregation.

In subsection 3.2.3, we combine these adjustments — varying both niche structure and percapita immigration rates across islands — to provide proof-of-principle that a niche–neutral model can reproduce the observed occurrence patterns. The final combined model is intentionally overfitted; its purpose is to demonstrate that the identified mechanisms are jointly sufficient, not to estimate parameter values.

#### 3.2.1. Allowing per-capita immigration rates to vary across islands

When we used the *θ* and *n*^∗^ parameter values from the homogeneous model fit and tuned the value of *m*_*i*_ for each island *i* to match observed species richness, we found that this brought the nestedness results into closer agreement with the survey data (Fig. 5). The observed raw nestedness remained lower than the model (data raw NODF 57.2; model raw NODF 70.2, 95% PI [63.3, 76.2], *p <* 0.01) but the standardised nestedness was within the range of the model (data SES NODF -2.67, model SES NODF −1.57, 95% PI [−4.45, 0.653], *p* = 0.19). Meanwhile, the data remained segregated compared to the model (model mean raw C-score 0.818, 95% PI [0.498, 1.18], *p* < 0.025; mean SES C-score: −0.753, 95% PI [−1.74, 0.660], *p* < 0.025).

**Figure 5.**
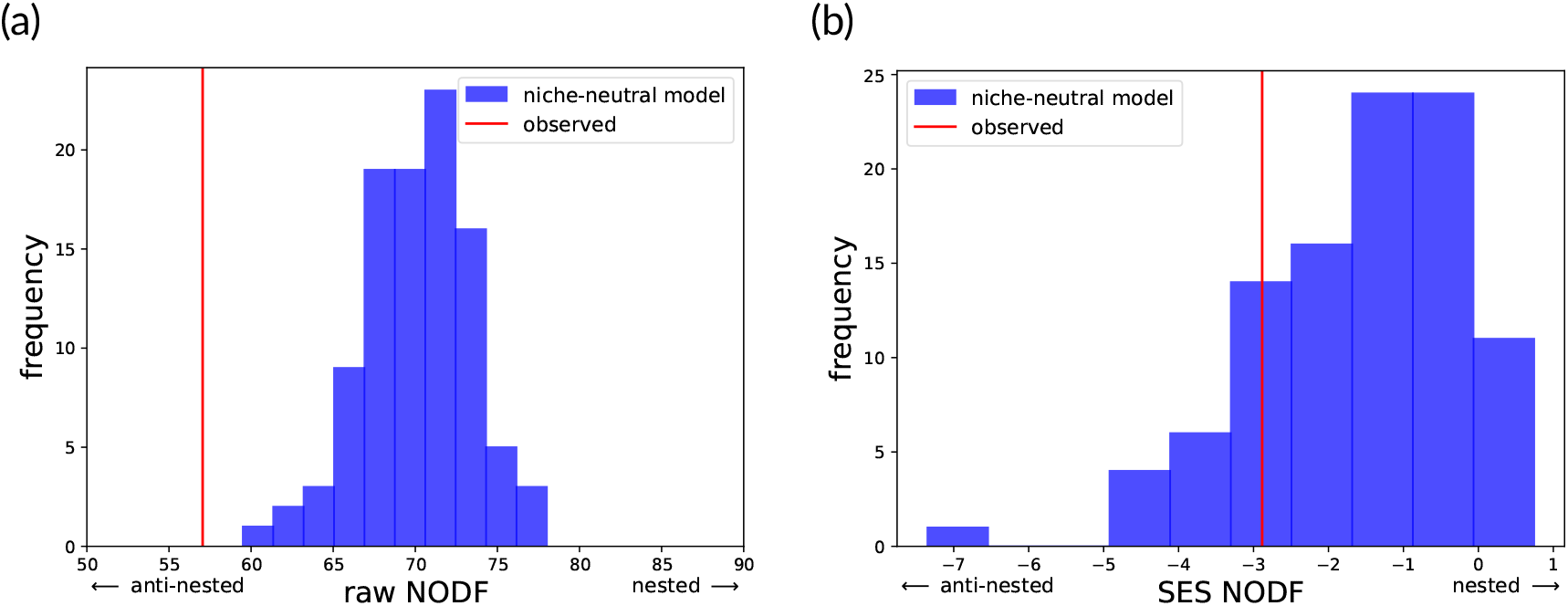
Frequency histograms comparing nestedness metrics between the observed data (red) and fitted niche–neutral models (100 draws, blue): (a) Raw NODF; (b) SES NODF. The niche–neutral model was fitted by allowing immigration rate *m* to vary to match the observed species richness on each island. The SES is calculated in comparison to fixed–fixed randomised matrices, generated using the sim9 algorithm with an effective sample size of at least 1000.

#### 3.2.2. Allowing niche structure to vary with island area

The main effect of allowing niche structure to vary with island area was to increase segregation (Fig. 6), with a smaller secondary effect of decreasing nestedness (additional results in SI E).

**Figure 6.**
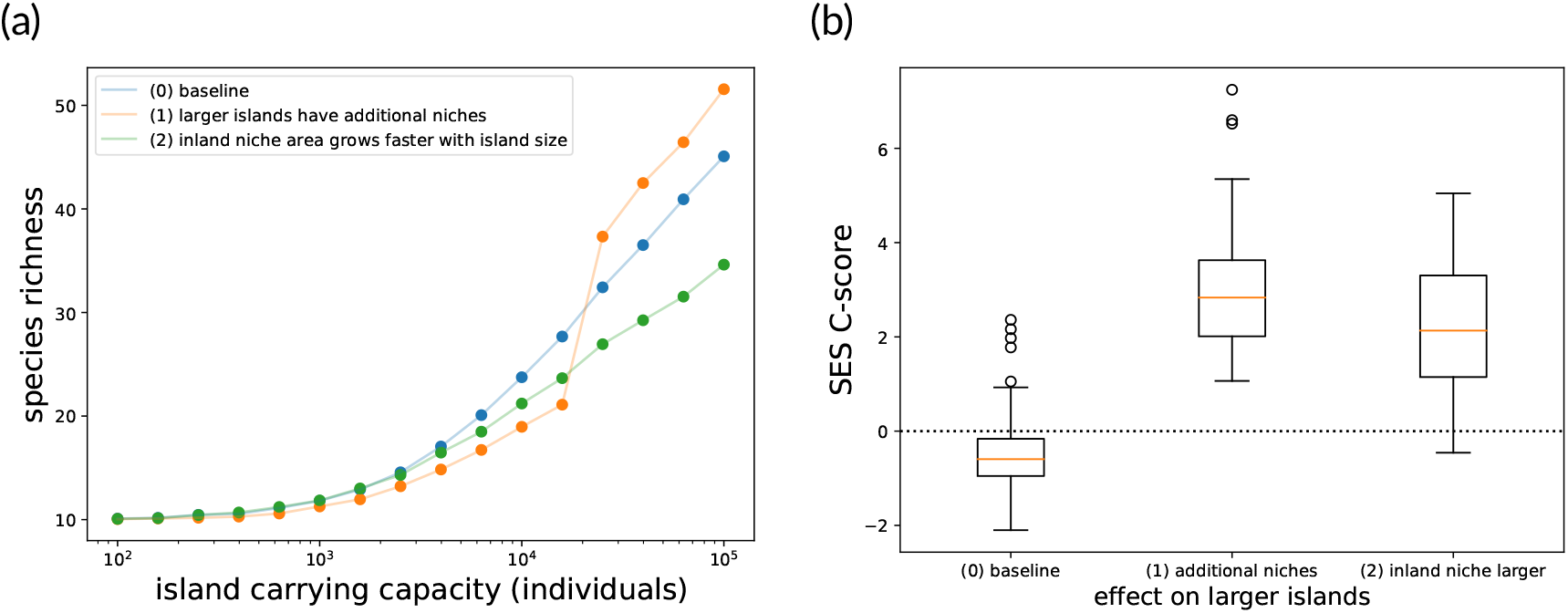
Effects of changing niche diversity with island area under three scenarios. In the baseline scenario (0), the homogeneous scenario, each island has *n*^∗^ = 10 equalarea niches and niche areas grow proportionally with island area. In scenario (1), the four largest islands have eight additional niches each. In scenario (2), the ten niches are split into coastal and inland types, and the area of coastal niches grows more slowly with island area. (a) The relationship between species richness and carrying capacity, which is a proxy for island area, under the three scenarios. (b) The SES C-scores under the three scenarios. Model parameters were *θ* = 10, *m* = 7.5 × 10^−4^. The results for each scenario represent 100 random model realisations using the sequential sampling algorithm.

A homogeneous model, with constant *m* and *n*^∗^ across islands, was used as a baseline to compare scenarios. The baseline model had the following occurrence-metric values: mean raw C-score 2.09 (95% PI [1.34, 2.80]), SES C-score −0.506 (95% PI [−1.84, 1.88]), raw NODF 75.6 (95% PI [69.5, 81.4]), and SES NODF −1.32 (95% PI [−3.73, 0.730]).

Scenario 1 (orange line, Fig. 6a) increased segregation compared to the baseline when using standardised scores (mean raw C-score 1.31, 95% PI [0.797, 1.94]; mean SES C-score 2.94, 95% PI [1.28, 5.96]) (Fig. 6b) and had a small negative effect on raw nestedness (mean raw NODF 74.3, 95% PI [69.9, 78.4]) and anti-nestedness (mean SES NODF −3.40, 95% PI [−5.98, −0.499]). Qualitatively similar results were found in other scenarios (online repository). To verify that the segregation effect in Scenario 1 was not simply because of the increased richness on larger islands, we modified the homogeneous model by fitting the immigration parameters (*m*) individually to larger islands to match the increase in richness in Scenario 1. This increased segregation, but not by as much (mean raw C-score 0.507, 95% PI [0.263, 0.788], mean SES C-score 0.269, 95% PI [−1.56, 2.14]).

Scenario 2 (green line Fig. 6a; details in SI D) also increased segregation compared to the baseline homogeneous model (mean raw C-score 3.77, 95% PI [2.51, 5.19]; mean SES C-score 2.25, 95% PI [−0.150, 4.88]) (Fig. 6b). Nestedness decreased and thus the archipelago remained anti-nested according to the standardised scores (mean raw NODF 64.0, 95% PI [58.3, 69.7]; mean SES NODF −1.68, 95% PI [−3.84, 0.900]).

#### 3.2.3. Varying both niche diversity and immigration rates across islands

Having verified that varying niche diversity across islands increases segregation (subsection 3.2.2), and that tuning immigration rates to match the richness data decreases nestedness 3.2.1), we combined these two observations to test if a modified niche–neutral model could plausibly reproduce the observed nestedness and segregation patterns. We assumed the number of niches increased with island size, with some niches occurring exclusively on large islands, and we tuned island-specific parameters (*n*^∗^ on small islands and *m* on large islands) to match the species richness pattern across islands. Through manual calibration, we found parameter values (details in SI F) that brought the niche– neutral model’s segregation and nestedness patterns into closer agreement with the survey data (mean SES C-score 6.16, 95% PI [4.02, 8.80]; mean SES NODF −4.39, 95% PI [−7.78, −1.49]; Fig. 7; Table 1 “Fitted”).

**Figure 7.**
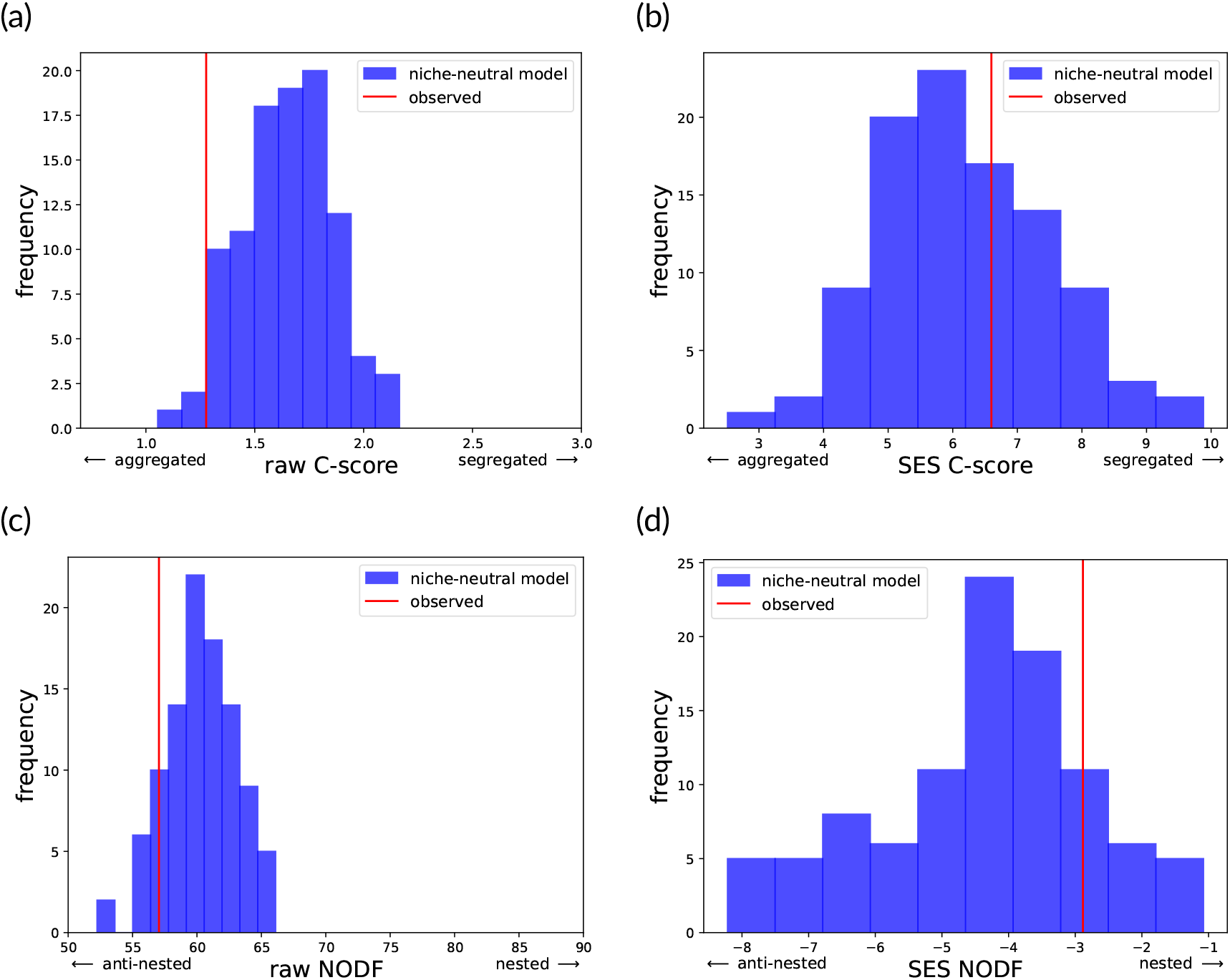
Frequency histograms comparing the distribution of occurrence characteristics of the observed survey data (red) to a niche–neutral model with niche diversity and immigration parameters varying across islands and parameters tuned to match the data (100 draws, blue): (a) raw C-score, (b) SES C-score, (c) raw NODF, (d) SES NODF. Details are provided in SI F.

## 4. Discussion

We used a sequential sampling algorithm to efficiently explore species occurrence patterns in niche–neutral models. A niche–neutral model serves as a null reference by representing the system with particular ecological mechanisms of interest excluded; specifically, species that share the same niche behave neutrally, which removes competitive-exclusion effects. This mechanistic approach was able to identify patterns of segregation and nestedness in community data, similar to traditional data-randomisation approaches, and the homogeneous niche–neutral model was rejected as a null model for bird species-occurrence patterns in Riau. However, when predictions do not match observations, this modelling approach offers a distinct advantage: it provides a framework for incorporating additional mechanisms into refined models. In our case-study, where richness is governed by immigration rate and the number of non-overlapping, internally neutral niches, we were able to verify that spatial heterogeneity in these parameters could plausibly reproduce the observed patterns. In other words, the anti-nestedness and segregation observed can be explained by niche–neutral processes, without the need to invoke functional differences between species within the same niche. Alternative modelling frameworks could explore different mechanisms (e.g., trait-based environmental filtering in the ecolottery model, Munoz et al., 2018). While such explorations cannot definitively identify *the* underlying mechanism, they can reveal that patterns previously attributed to one process (e.g., segregation due to competition) may arise through alternative pathways (e.g., higher niche diversity on larger islands).

Sequential sampling algorithms offer a significant advancement in the efficiency and application of neutral models for studying occurrence patterns (Munoz et al., 2018). Neutral models have been valued in previous studies for their parsimony, simplicity, and mechanistic basis: they rely on minimal assumptions, are based on first principles, and yield rich predictions with few parameters by assuming functional equivalence between species (Marquet et al., 2014). However, previous applications to co-occurrence analysis have relied on computationally heavy forwards-in-time simulations (Bell, 2005; Gotelli and McGill, 2006; Ulrich, 2004; Ulrich et al., 2017). Sequential sampling algorithms facilitate more efficient applications by massively accelerating simulations (Munoz et al., 2008, 2007, 2018, 2014). The algorithm is based on the coalescence approach to neutral modelling (Rosindell et al., 2008), which involves sampling the genealogy of a present community backwards in time. This backward-in-time approach relies on the per-capita equivalence of species within niches, an assumption that allows for efficient simulation. Therefore, a prudent general approach to analysing species occurrence patterns may be to start with a computationally efficient neutral or niche–neutral model, and once key missing mechanisms have been identified, move on to more tailored, albeit likely more computationally expensive, models (e.g., see a series of papers disentangling species distribution patterns in ground beetles: Ulrich and Zalewski (2006, 2007) and Zalewskia and Ulrich (2009)).

In our application to birds in Riau, we demonstrated how starting with a simple niche–neutral model and then increasing its realism identified potential mechanisms behind species occurrence patterns. The simplest version of the niche–neutral model—the homogeneous model—could fit the species–area relationship, but it failed to accurately describe total species richness and occurrence patterns. Specifically, the data showed greater richness, greater segregation and anti-nestedness compared to the model. A traditional data-randomisation analysis lead to qualitatively similar results (Table 2), but the advantage of the mechanistic approach is that it also provided a quantitative framework for testing hypotheses about missing mechanisms. In our case study, we showed that by relaxing the homogeneity assumption and increasing the model’s realism, the model better matched observations. Specifically, species segregation increased when we allowed habitat diversity to increase with island size, and nestedness decreased when we tuned model parameters to more accurately reproduce the richness pattern across islands (detailed below). Thus, we conclude that differences in habitat diversity and per-capita immigration rates across islands, while not necessary for explaining coarse patterns like the island species– area relationship, may be crucial for explaining species-occurrence patterns.

Varying niche diversity across islands increased species segregation because islands with different niches (scenario (1) in Fig. 6) or different-sized niches (scenario (2)) tend to have fewer species in common than those with homogeneous niches (scenario (0)). Previous studies have noted that the positive effect of island area on species richness may be partially mediated by habitat diversity (Kohn and Walsh, 1994), particularly if large islands contain key habitats (e.g., rainforests, Davidar et al., 2001) and the species group under consideration includes habitat specialists (Ricklefs and Lovette, 1999). Scenario (2) has the advantage of parsimony: it simply adds the biologically reasonable assumption that coastal niches grow linearly with island perimeter, while other niches grow linearly with island area.

Varying per-capita immigration rates across islands decreased nestedness because it disrupted the relationship between island size and species occurrence. Nestedness has been attributed to several mechanisms, including selective immigration, passive sampling, and stochastic colonisation-extinction dynamics (Higgins et al., 2006; Lomolino, 1996; Wright et al., 1998). In the homogeneous model, species on smaller islands are typically subsets of larger islands because smaller islands have both lower total immigration rates (per-capita immigration rate times area) and higher extinction rates (MacArthur and Wilson, 1967). Choosing immigration rates that best fit the island-richness pattern weakens the dependence of occurrence patterns on area and thus reduces nestedness. A limitation of our exploration of this phenomenon is that we allowed the per-capita immigration rate for each island to vary independently, leading to an over-fitted model. Therefore, our analysis should be viewed as simply proof-of-principle that accounting for inter-island variation in richness, such as by varying per-capita immigration rates, reduces nestedness.

Readers should exercise caution when interpreting the fundamental biodiversity estimate *θ* obtained in this study. Technically, *θ* represents the rate at which new species arise in the metacommunity through speciation. However, the regional pool from which immigrants arrive to local communities may itself be a subregion partially assembled by immigration from a larger metacommunity (Munoz et al., 2008). In such cases, *θ* represents a combination of regional speciation plus immigration from the larger-scale metacommunity.

Further work is needed to identify the niche processes operating in Riau, which may require going beyond the simple non-overlapping niche description of the niche–neutral model. The niche–neutral model assumes *n*^∗^ discrete, non-overlapping niches that govern the niche-structured dynamics. However, the actual niches of birds (and other taxonomic groups) heavily overlap, and previous authors have noted that it is unclear what is being represented by the “habitats” that influence island species diversity (Triantis et al., 2003, 2006). This suggests that the *n*^∗^ parameter in our model may be better thought of as an effective parameter that describes how habitat diversity affects species diversity (cf., Triantis et al., 2003, 2006).

A more detailed analysis could also consider the mechanistic basis for variation in immigration rates, which we conjecture is mainly due to differences in island isolation. For example, Durian and Ngenang, although smaller than Temiang and Sebangka (Fig. F.2), have higher species richness because they are closer to the large land masses of Sumatra and continental Asia, respectively, and larger nearby islands may also play a role. Quantifying island isolation and incorporating it into a per-capita immigration rate for each island would be a more parsimonious way to explore this phenomenon; however, that would also require more data and increase the complexity of the fitting process. In particular, if a niche–neutral model is used as a starting point, and later improved to account for more complex immigration (or niche, above) processes, it is likely that the parameter values of the initial model will be biased and not provide a correct assessment of the contribution of neutral processes (Gotelli and McGill, 2006).

Our exploration of these scenarios is intended as illustrative of a general idea: when a mechanistic neutral or niche–neutral model is used, it can not only serve as a null model, but also be extended to identify other potential mechanisms driving occurrence patterns. To be explicit about the epistemological status of our fitted model: the island-specific parameter values are not intended as estimates. The model is overfitted by design, because its purpose is diagnostic. The question we asked was not “is this the right model?” but “can the mismatches between the homogeneous null and the data be resolved by mechanisms that are ecologically plausible within the niche–neutral framework?” The answer — that varying habitat diversity and immigration rates across islands is jointly sufficient — identifies these as candidate mechanisms worth investigating with independent data, not as confirmed explanations. Our analysis is intended to demonstrate the diagnostic value of the approach itself.

Future workers with more data may wish to apply a more systematic model-fitting approach than the method we used here. For example, if full species abundance distribution data is available, then the R package `ecolottery`can be used to simulate neutral dynamics (Munoz et al., 2018), and it includes Approximate Bayesian Computation (ABC) tools (Sisson et al., 2018) for parameter estimation. These tools can be used to estimate immigration rates and environmental filtering parameters, compare observed and simulated community patterns, and test hypotheses about community assembly mechanisms (e.g., Blanchard et al., 2020; Denelle et al., 2019). However, if one wishes to investigate the nestedness and co-occurrence metrics we used, then the computational effort may be prohibitive. Specifically, although obtaining a single sample from the sequential sampling algorithm is relatively fast (Fig. 2c-d), one must randomise each sample a large number of times to obtain the SES metric recommended in the literature (Fig. 2e), which is computationally expensive. The species-occurrence literature recommends the fixed-fixed randomisation (Gotelli and McCabe, 2002; Ulrich et al., 2018), which requires the use of the curveball algorithm (Strona et al., 2014), which produces autocorrelated samples. Therefore, the curveball algorithm must be run for a large number of iterations to obtain a sufficiently large effective sample size. Alternatively, a fixed row-or column-sum randomisation may be feasible for certain applications. For example, the PER-SIMPER tool has been used to distinguish niche-from dispersal-dominated community assembly (Gibert and Escarguel, 2019), which can be summarised in a single metric (DNCI, Vilmi et al., 2021) that might be used in a similar way to SES co-occurrence and nestedness here.

A distinct advantage of our approach is that it only requires species presence-absence data for each island, rather than the full species abundance distribution (SAD) data typically needed to parameterise neutral models (Beeravolu et al., 2009). However, for future workers who have access to species-abundance data, we wish to highlight alternative sequential sampling algorithms in the literature. Munoz et al. (2007) provides a method that can estimate the immigration parameter for each island separately, in contrast to our default niche–neutral model (homogeneous model), which assumes a constant immigration rate across islands. Munoz et al. (2008) present a three-level spatially implicit neutral model comprising: a metacommunity where new species originate through speciation; a regional pool that receives immigrants from the metacommunity; and multiple local communities that receive immigrants from the regional pool. Crucially, by using Simpson-based similarity statistics to calculate their novel *G*_ST_(*k*) for each local-community sample *k* (analogous to the *G*_*ST*_ statistic from population genetics), they demonstrate that the immigration parameter for each local community can be estimated without requiring either an estimate of the metacommunity *θ* nor assumptions about the species abundance distribution of the regional pool, which may match the metacommunity (as in Hubbell’s two-level model) or deviate from it. Munoz et al. (2014) builds on Munoz et al. (2007) and introduces explicit modelling of habitat filtering (similar to our *n*^∗^ distinct, non-overlapping niches) with the option of varying niche conservatism (our model assumes perfect niche conservatism). The models in Munoz et al. (2008) and Munoz et al. (2014) were brought together in an R package called `ecolottery`(Munoz et al., 2018), which includes the option to model habitat-filtering effects, which may be useful for future workers with species-trait data. For example, by fitting the immigration parameter plus three environmental optimum parameters and their standard deviations to the top three principle components in species-trait space, Blanchard et al. (2020) were not only able to demonstrate that communities deviated from neutral expectations (the environmental filtering parameters were necessary), but also that environmental filtering targets different traits differently (e.g., near fragment edges, there is strong selection for resource acquisitive species) (see also, Denelle et al., 2019). For workers interested in the question of how environmental filtering and niche separation have affected community assembly, a summary statistic measuring deviation in functional diversity from neutral theory is available (*u* statistic, Laroche et al., 2020).

Previous workers have suggested using neutral models as the null model for species occurrence analysis; however, we expect that, in most cases, neutral models will be ruled out as exact models due to mismatches with the species-richness pattern. In our case study, although the niche–neutral model could explain the overall trend in the island species–area relationship, it exhibited insufficient variability to explain the individual observed island richnesses (Fig. 3a). Therefore, its failure as an exact descriptor is clear even before considering more detailed metrics such as segregation and nestedness. We expect this situation will be common (compare with Louca et al. (2016) and random-placement models below), and the real value of neutral models is as an exploratory framework as described above, for exploring alternative ‘how-possibly’ explanations (Zhang, 2020).

Previous work with various neutral-style models have observed a similar lack of variability, which ruled them out before considering more detailed metrics. For example, Louca et al. (2016) fitted a neutral model designed for microbial systems. Their model reproduced the positive relationship between abundance and detection frequency, analogous to our positive relationship between area and species richness. However, they found that a large proportion of their taxonomic groups fell outside the probable range of model predictions, just as we found that 16 of 23 observed island richnesses fell outside the model’s 95% prediction intervals (Fig. 3a). Analogous results are also obtained from random-placement models, which simulate passive sampling by assuming that the probability of a species’ presence at a site depends only on its relative regional abundance (Coleman, 1981; Coleman et al., 1982; Dolman and Blackburn, 2004). These models are known to produce nested patterns independent of other ecological mechanisms (Cutler, 1994; Wright et al., 1998), and therefore they are often used as a preliminary test before more detailed nestedness analyses (Worthen et al., 1998). However, workers often find that the observed site species richnesses fall outside the bounds predicted by a random-placement model, which rules out passive sampling as an explanation for the data (e.g., Calmé and Desrochers, 1999; González-Oreja et al., 2012; Tan et al., 2021; Wang et al., 2010). Our niche–neutral model retains most of the assumptions of neutral models, and thus, these previous studies and our results tell a consistent story: while neutral models and their ilk may reproduce coarse biodiversity patterns, their predictions lack the variability of empirical data.

We suggest three general reasons why neutral models predict less variability than empirical data. First, these models are spatially implicit. They cannot account for how species composition changes over space due to dispersal limitation and the decline of relatedness with distance (Rosindell and Cornell, 2007), which also leads to less accurate predictions of macroecological patterns compared to spatially explicit models (Rosindell and Cornell, 2009). Second, these models typically ignore spatial variability in model parameters (c.f. Dolman and Blackburn, 2004) as shown by the improved fit to occurrence patterns when we allowed niche diversity and percapita immigration rates to vary with island area (Figs. 5–7). Finally, these models ignore temporal variability in model parameters, which in nature is important because of temporal environmental stochasticity (Fung et al., 2016). In contrast, the only type of stochasticity in neutral models and our niche–neutral model is demographic stochasticity, which leads to much smaller fluctuations in species abundances over time than does environmental stochasticity, particularly for abundant species. For these reasons, we expect that any future applications of our niche–neutral null model approach to analysing occurrence patterns will also likely reveal higher-than-expected segregation and lower nestedness, as found here.

We conclude that mechanistic models such as the niche–neutral model offer a qualitative advance over data randomisation for analysing species occurrence patterns. Data randomisation can identify non-random patterns but provides no path forward when non-randomness is detected. A mechanistic approach, by contrast, turns rejection of the null into an iterative diagnostic process: each mismatch between model and data implicates a candidate mechanism, and modifications that resolve specific mismatches identify which processes are likely to matter. The sequential sampling scheme facilitates such applications by accelerating simulations of neutral and related models. To these ends, we hope that the code and tutorials we have provided for the algorithm (SI G) will be useful to future workers in this area.

## 5. Acknowledgements

We thank Tak Fung for helpful comments, and we thank F. Munoz, who served as Recommender for this manuscript, for detailed and constructive feedback across multiple rounds of review that substantially improved the clarity and framing of the paper.

A preprint version of this article has been peer-reviewed and recommended by PCI Ecology (https://doi.org/10.24072/pci.ecology.100770).

## 8. Data, script, code, and supplementary information availability

Code is available online https://github.com/nadiahpk/niche-neutral-riau-birds, archived at https://doi.org/10.5281/zenodo.19810402, and described in the Supplementary Information.

## Supplementary Information

**Using a sequential sampling algorithm to apply the niche-neutral model to species occurrence patterns**

N. P. Kristensen, Y. C. K. Sin, H. S. Lim, F. E. Rheindt, R. A. Chisholm

## Appendix A. Sequential sampling scheme algorithm for the niche–neutral model

Algorithm 1 provides an overview of the sequential sampling scheme. Tutorials and the implementation are available in the Github repository (nadiahpk/niche-neutral-riau-birds/):

1. `./tutorial/Sequential_Sampling.pdf`, `./tutorial/Sequential_Sampling.ipynb`: Tutorials explaining how the algorithm works using a concrete example.
2. `draw_sample_species_generator_general()`in `./functions/my_functions.py`: A function implementing the algorithm, which was used to generate results for this study.

### Algorithm 1

Sequential sampling scheme for the niche–neutral model

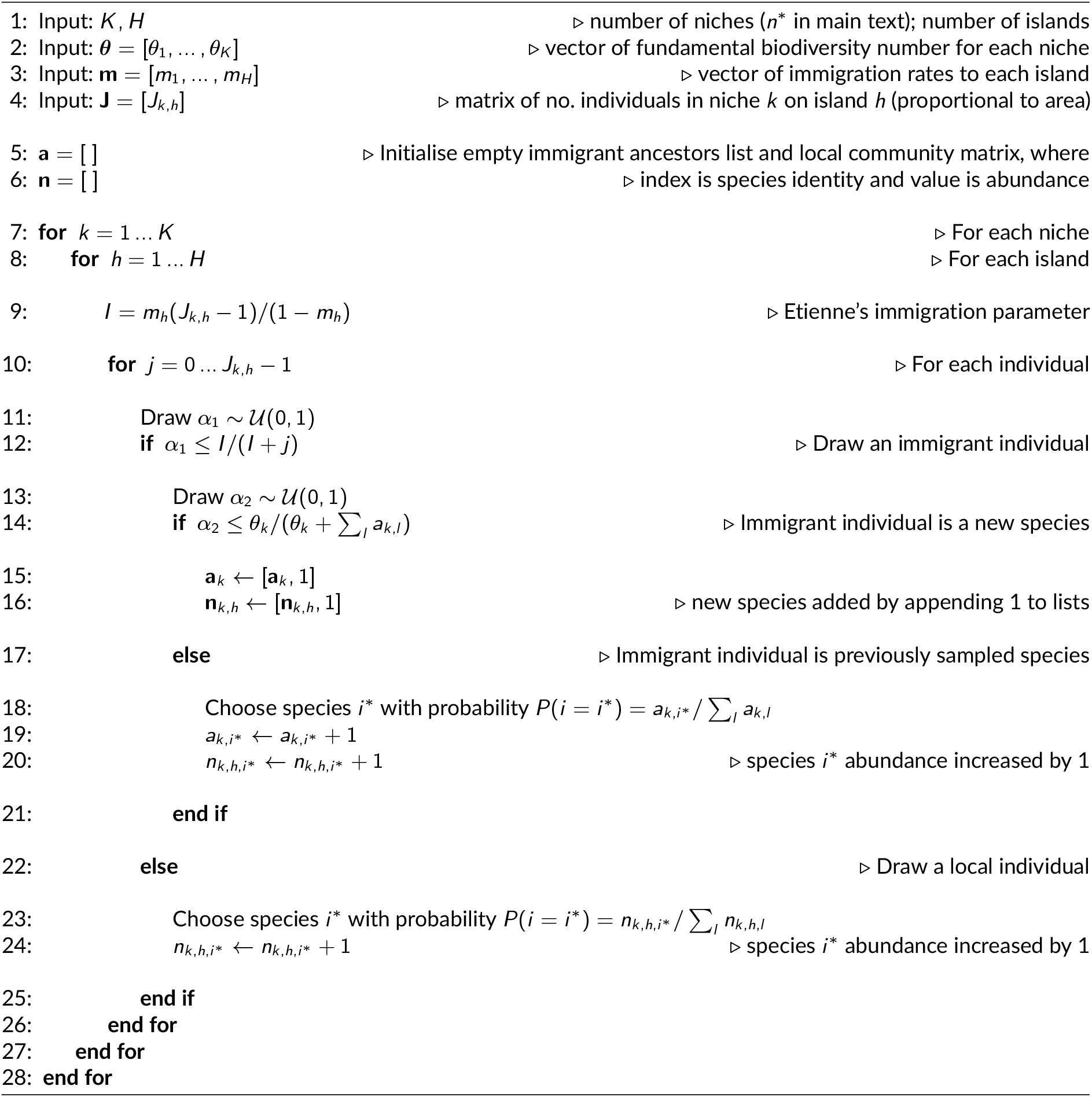

## Appendix B Fitting the homogeneous niche–neutral model

To parameterise the niche–neutral model, three model parameters — number of niches*K* (*n*^∗^ in main text), immigration rate *m*, and fundamental biodiversity number *θ* — were fitted to the island species–area data using the methods described in Chisholm et al. (2016). The fitting function is `fit_area_richness()`in the code repository. The notation used in this appendix is listed in Table B.1.

**Table B.1.**
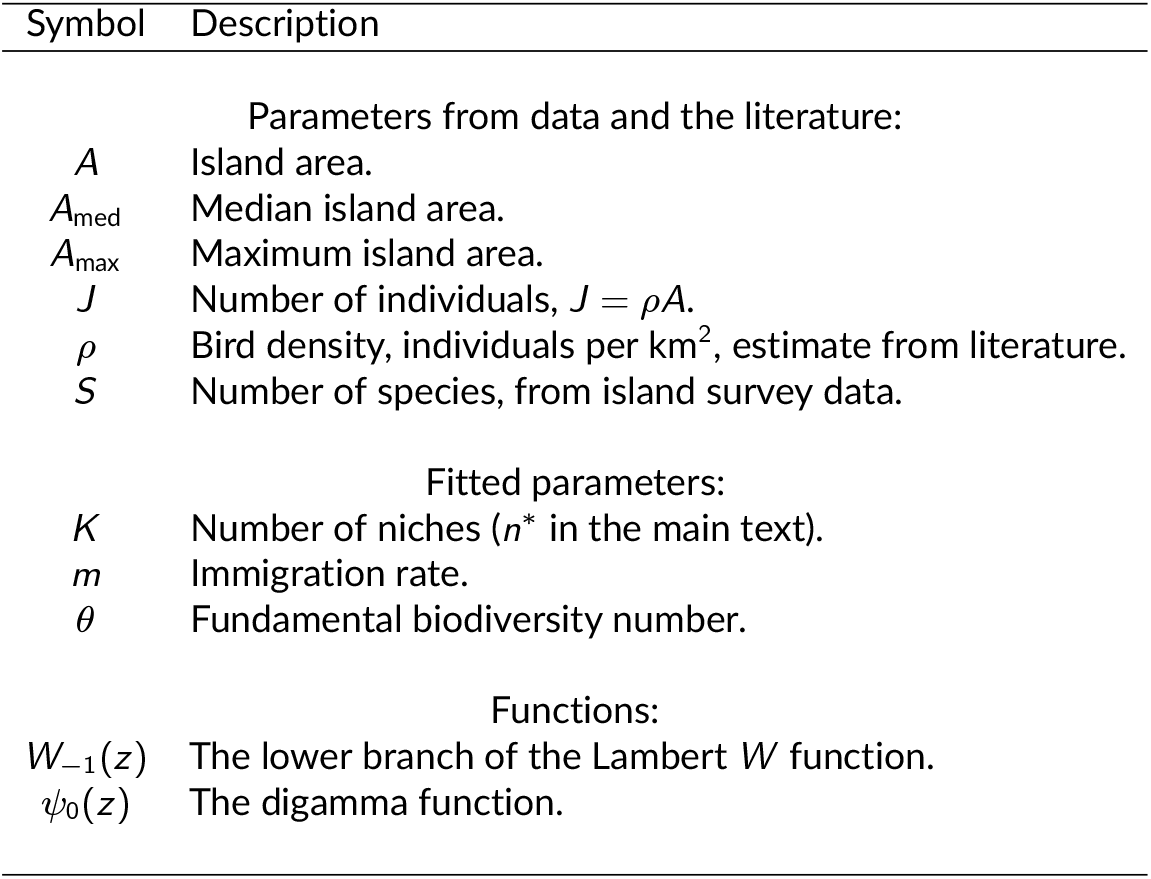
Notation.

In overview, the procedure is to find the best-fit values of *m* and *θ* for each value of *K*, and then choose from among the *K* the (*K, m, θ*) triple that gives the highest *R*^2^ value.

The species richness function that is fitted to the species-area relationship is (Eq. 2.5 Chisholm et al., 2016)

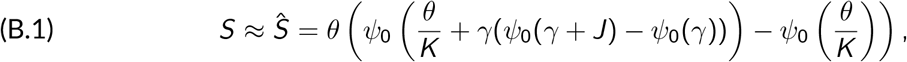

where *γ* = (*J* − 1)*m/*(1 − *m*), and the best-fit *m* and *θ* values were found by minimising the residual 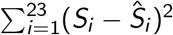, where the islands have been indexed *i* = 1, …, 23.

We found the best-fitting (*m, θ*) pair for each value of *K* ∈ {1, 2, …, 15}.

For *K* = 1, the initial guesses for *m* and *θ* were (S. 2(c) Chisholm et al., 2016)

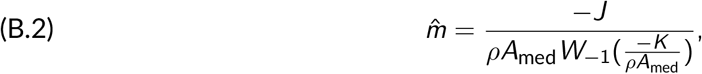

and

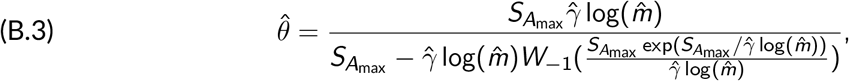

where 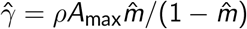.

For *K* ∈ {2, 3, …, 15}, we used the best-fitting values from the previous *K* as the initial guesses for *m* and *θ*.

## Appendix C Presence-absence matrices

The presence-absence matrices provide an intuitively visual way to compare the observed species distribution to the various model predictions (Fig. C.1).

Comparing the observed matrix (Fig. C.1a) to examples of predictions from the homogeneous niche–neutral model (Fig. C.1b, c), it is apparent that the model predicts lower total richness and higher nestedness than observed.

When both niche diversity and immigration rates were varied across islands (S 3.2.3), matrices were visually similar to the observed matrix (Fig. C.1d-e).

**Figure C.1.**
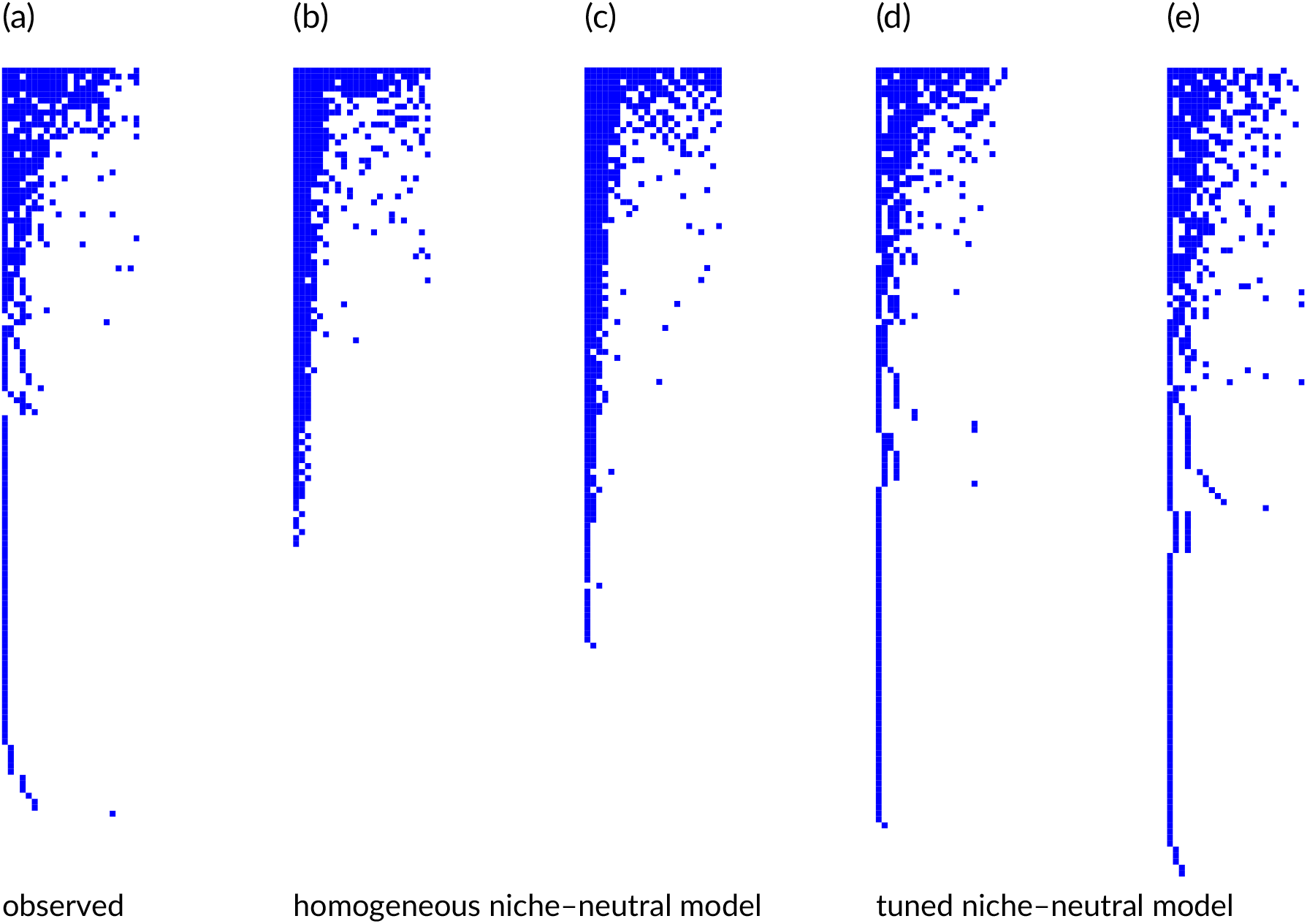
Comparing presence-absence matrices from (a) observations, (b-c) bestfitting homogeneous niche–neutral model, (d-e) scenario contrived to match segregation and nestedness. Rows are species, columns are islands, and a coloured square represents the presence of a species on an island.

## Appendix D Details of Scenario 2 (width of coastal niches held constant)

In Section 3.2.2, we explored if allowing the niche structure to vary with island area would increase segregation in the niche–neutral model. In Scenario 2, we explored a scenario where the width of coastal niches around the perimeter of the island (assumed circular) remained constant across the archipelago (Fig. D.2). The purpose of this appendix is to detail how this was done.

**Figure D.2.**
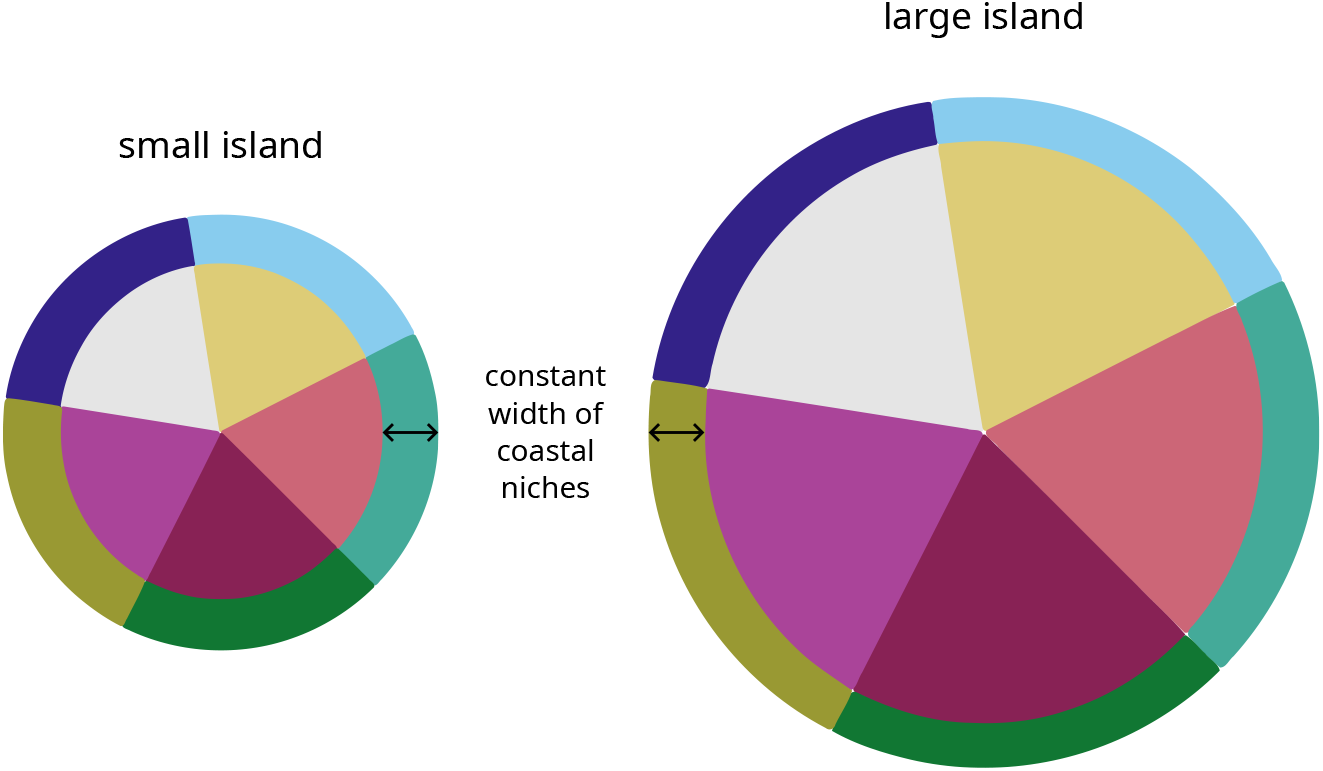
An illustration of Scenario 2, where each coloured segment represents the proportion of the island covered by each niche. Niches are split into two types — inland and coastal niches — and the width of the coastal niche area around the perimeter of the island (assumed circular) is held constant as the size of islands in the archipelago varies.

We fixed the width of the coastal niches *w*_*C*_ such that the area of coastal niches *A*_*C*_ was equal to the area of inland niches *A*_*I*_ when the total area *A* = *A*^∗^. Then the fixed width of coastal niches was calculated as the difference between the total radius and the radius of inland niches when 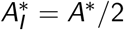:

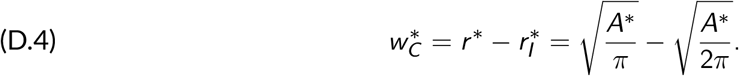

For other islands in the archipelago with area *A* with radius *r*, the areas covered by the inland niches *A*_*I*_ and coastal niches *A*_*C*_ were calculated by:

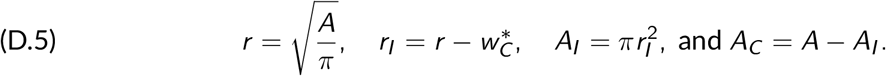

For these synthetic scenarios, we found it convenient to work directly in terms of numbers of individuals *J*. Recall that the number of individuals in an area is a linear function of the area, *J* = *ρA*. Therefore, the calculations above can be performed directly in terms of *J* by applying a transformation 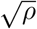 to distance measures. In other words, we define a ‘radius’ and the fixed coastal-niche ‘width’

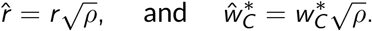

Then the fixed ‘width’ of coastal niches is calculated when 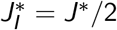

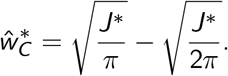

We explored the range of island carrying capacities *J* ∈ [10^2^, 10^5^], and for each new *J*, the number of individuals residing in inland and coastal niches was calculated:

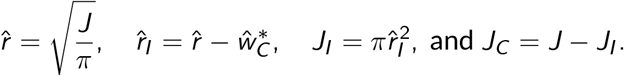

For Scenario 2, we set *J*^∗^ = 100, and we split the ten niches of our synthetic archipelago into five coastal and five interior niches of equal size within their types (Fig. D.2). The code used to generate Scenario 2 can be found in the code repository:

`scripts/neutral_vary_JK/create_archipelago_params_2.py`.

## Appendix E Raw nestedness decreases with increasing niche diversity

To further explore the effect of niche diversity on species distribution patterns, the niche– neutral model was refitted to survey data at varying fixed *n*^∗^ levels. Then, the fitted parameter values were used to create idealised archipelagos of 16 islands each (Fig. E.3a).

**Figure E.3.**
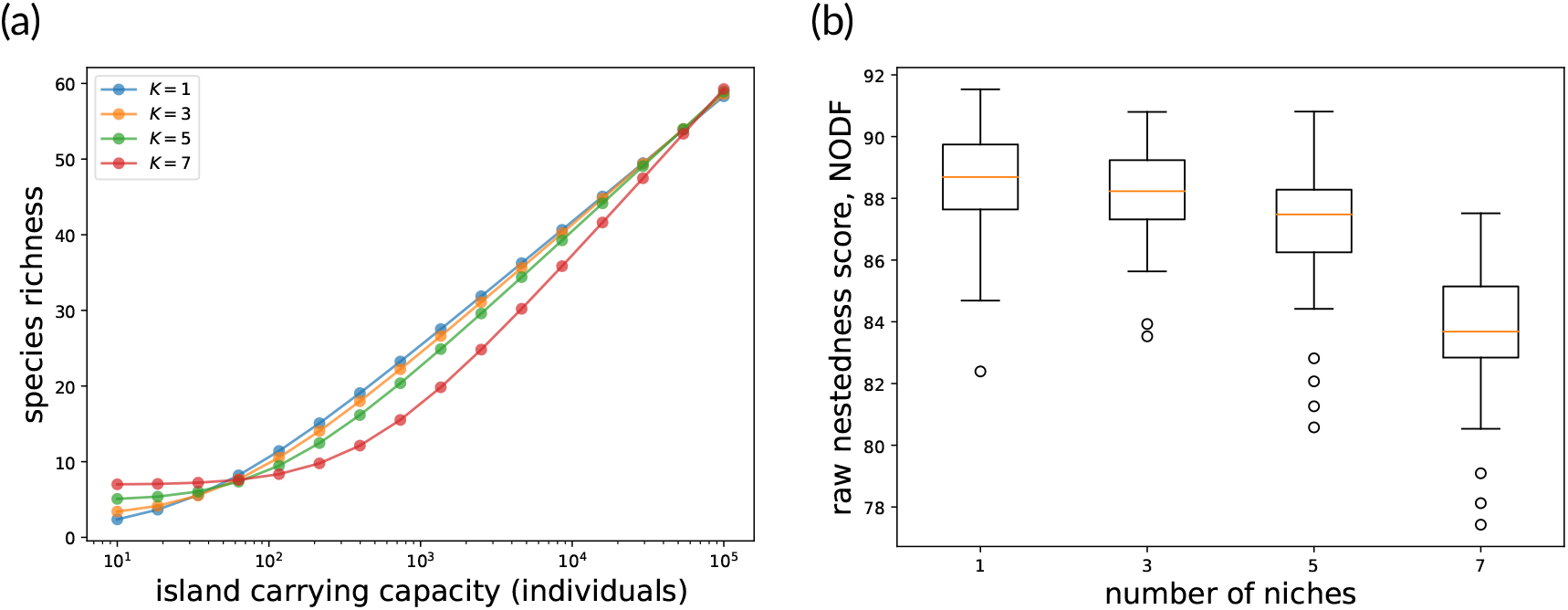
The effect of increasing the number of niches on every island on nestedness: (a) species richness versus carrying-capacity relationship; and (b) the effect of four *n*^∗^ scenarios on raw nestedness.

In general, increasing *n*^∗^ decreased the raw nestedness score (*n*^∗^ = 1 mean raw NODF 88.6, 95% PI [85.1, 91.5]; *n*^∗^ = 7 mean raw NODF 83.8, 95% PI [78.6, 86.6]; Fig. E.3b) but increased the nestedness compared to the fixed–fixed randomisation (*n*^∗^ = 1 mean SES NODF −1.49, 95% PI [−5.29, 0.91]; *n*^∗^ = 7 mean SES NODF −1.08, 95% PI [−4.18, 0.994]).

## Appendix F Details of the tuned niche–neutral model

In this appendix, we detail the example scenario presented in main text. The scenario was generated with the script: `scripts/neutral_data_fitmK/fitmK_3.py`. Additional experiments can be found in the electronic repository, in subdirectories named `*data_fit*`.

The observed data has a lower raw NODF and higher SES C-score than is produced by the best-fitting homogeneous niche–neutral model (Fig. 4). However, the homogeneity assumption, i.e., that all islands have the same *n*^∗^ and *m*, is unrealistic. In real archipelagos, habitat diversity typically increases with island size (Davidar et al., 2001; Kohn and Walsh, 1994; Ricklefs and Lovette, 1999), and the immigration rate varies in complicated ways depending on both the distance to the mainland and nearby islands as well as the species characteristics.

From the simulation experiments with hypothetical archipelagos (Sect. 3.2), we also learnt that (1) imposing a positive relationship between island size and number of niches *n*^∗^ increases SES C-score, and (2) tuning the island-specific immigration rates *m* so the model correctly predicts each island’s richness decreases raw nestedness. Therefore, we expected that combining these assumptions into a fitted niche–neutral model would bring the model predictions into closer agreement with observations.

The scenario presented in the main text was obtained through a combination of applying the 2 generalisations and a process of manual calibration. The total number of niches in the archipelago was increased from *K* = 8 (best-fit value in the homogeneous model) to 11. The extra niches represent niches that are available on some islands but not others, which increases species segregation. Any (small) island with fewer than 8 species was set with an island-specific *K*_*h*_ equal to the number of species. For islands with more than 8 species, the island-specific immigration parameter *m*_*h*_ was fitted so that the expected number of species on the island was equal to the observed richness. For some large islands, fitting *m*_*h*_ in this way led to unrealistically high immigration rates, several orders of magnitude greater than best-fitting rate from the homogeneous model (*m* = 0.0036). Therefore, those islands were instead given a higher island-specific niche diversity (*K*_*h*_ *>* 8), which is consistent with the observation that large islands can sustain key habitats like rainforests (Davidar et al., 2001), and which allowed a more realistic *m*_*h*_ to be fitted.

Table F.2 summarises the example scenario presented in the main text. Niches numbered 0 to 10 were distributed somewhat arbitrarily across islands, but low-numbered niches represent habitats that occur on most islands, and high-number niches (e.g., niches 9 and 10) represent habitats that typically occur only on larger islands (e.g., intact rainforest). Fig F.4 shows how the fitted *m* varied with island size.

**Table F.2.**
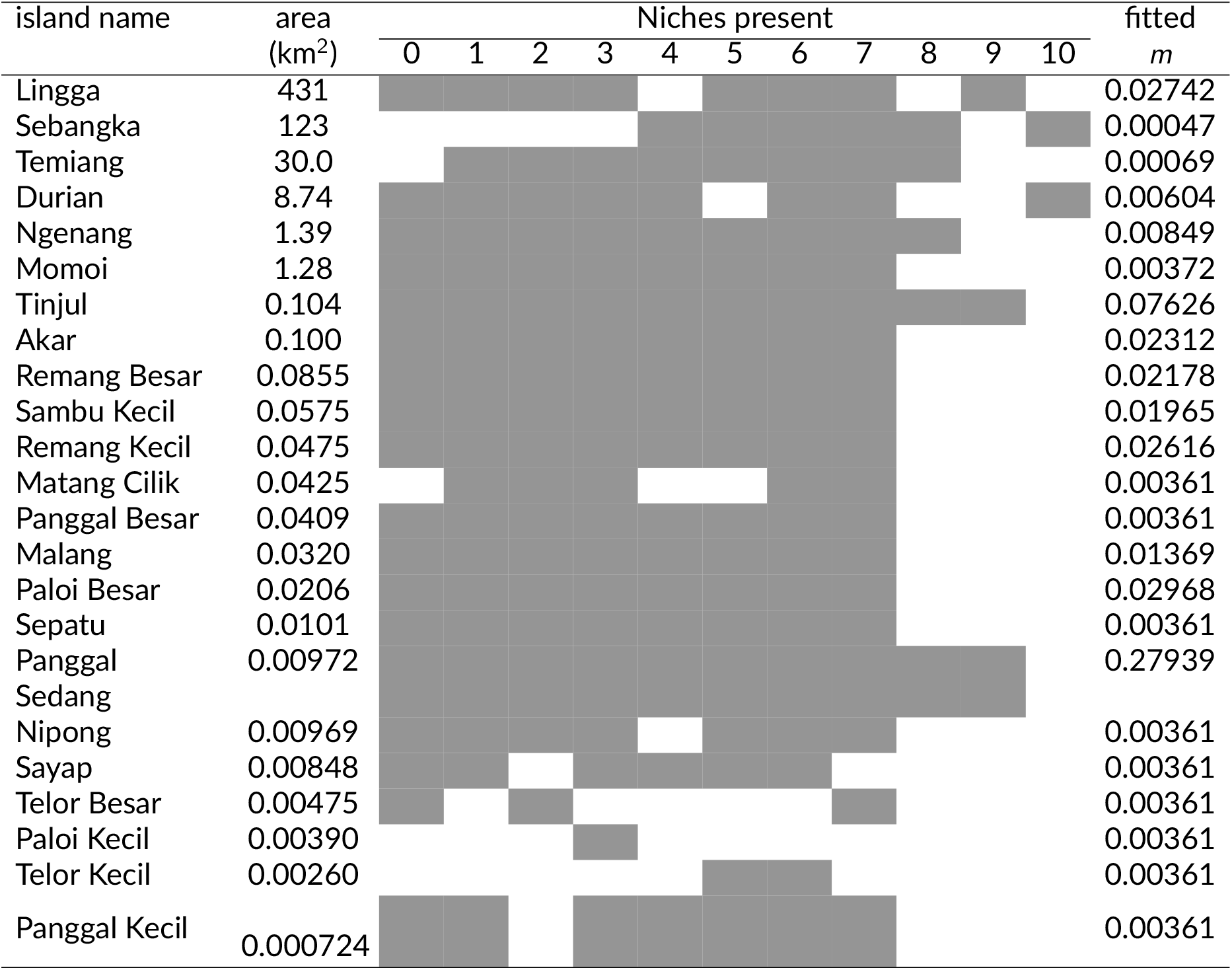
Summary of the tuned model presented in the main text.

**Figure F.4.**
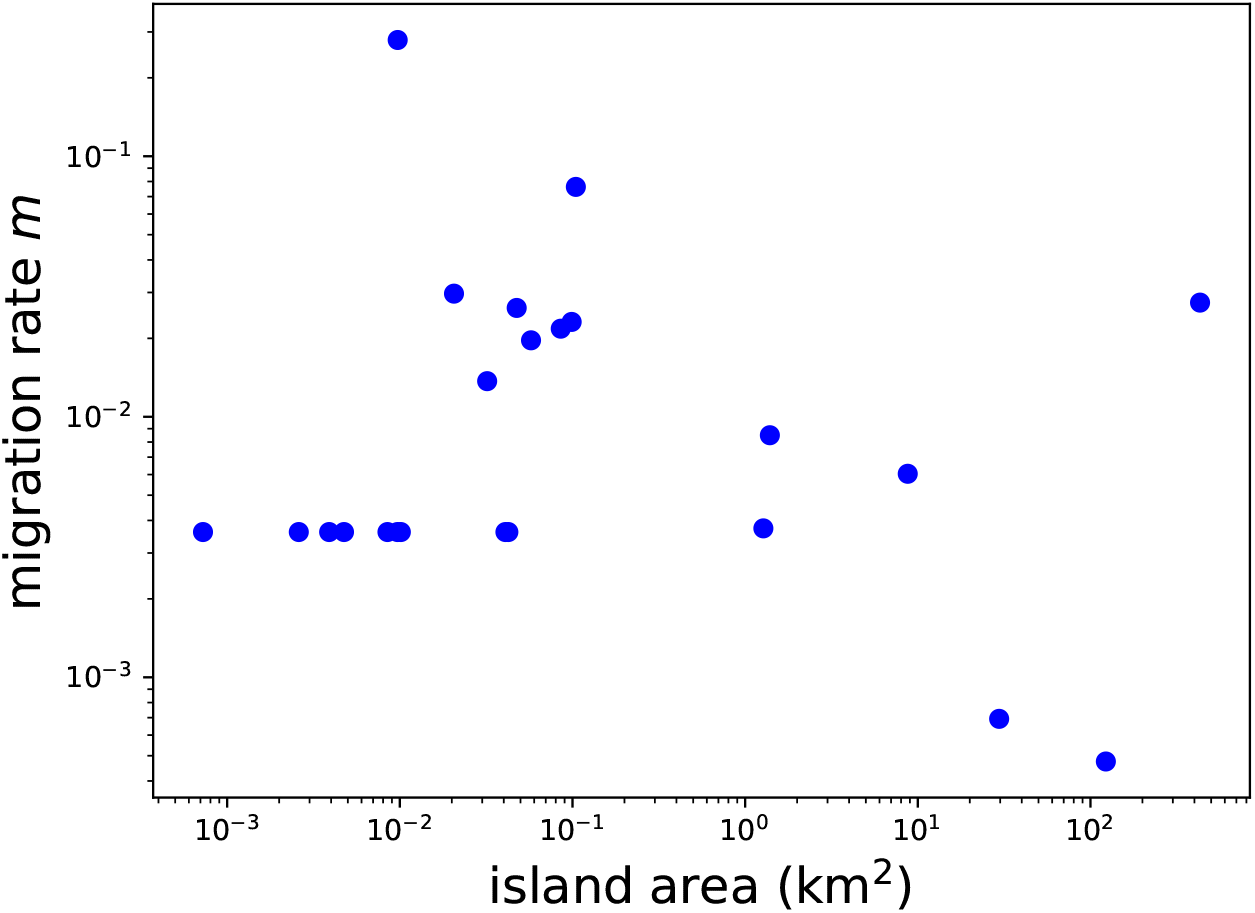
Fitted *m* versus island area.

## Appendix G Overview of the code repository

Functions and scripts used to generate the results are archived in the Github repository: https://github.com/nadiahpk/niche-neutral-riau-birds, archived at https://doi.org/10.5281/zenodo.19810402 (Fig. G.5).

**Figure G.5.**
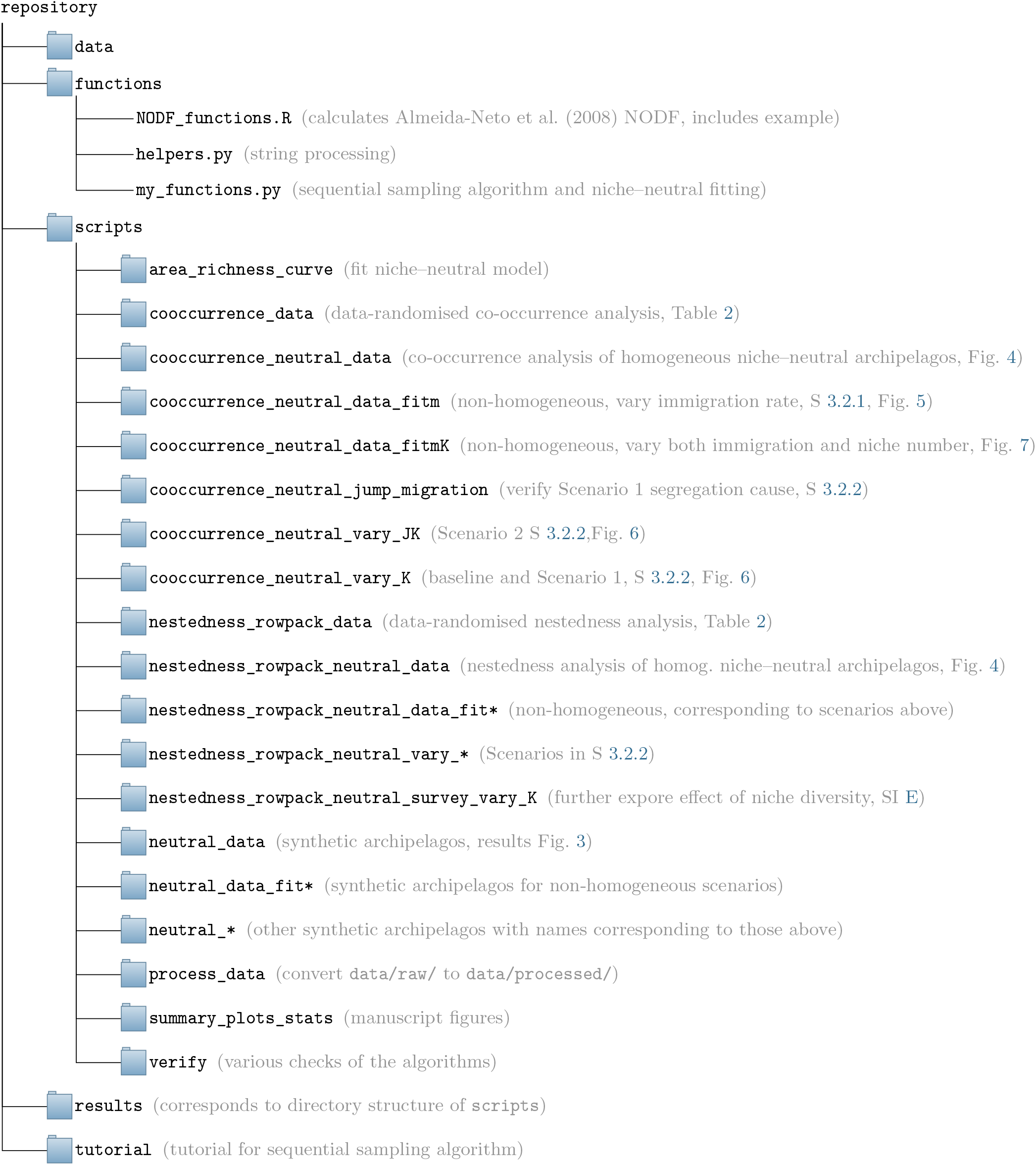
Overview of the contents of the Github repository.

C-score calculations are in scripts in directories `/scripts/cooccurrence*/`, and NODF calculations are in scripts in directories `/scripts/nestedness*/`.

Future workers who wish to use the niche–neutral model and sequential sampling algorithm may find the following resources useful:

(1) Tutorial for the sequential sampling algorithm: `/tutorial/`.

(2) Function implementing the algorithm is in `/functions/my_functions.py`: `draw_sample_species_generator_general()`and `draw_sample_species_generator()`.

(3) Function to fit the niche–neutral parameters to data is in: `scripts/area_richness_curve/fit_all_island_subsets.py`. It uses the function `fit_area_richness()`in `/functions/my_functions.py`.

